# *Entamoeba histolytica* develops resistance to complement deposition and lysis after acquisition of human complement regulatory proteins through trogocytosis

**DOI:** 10.1101/2021.08.04.455151

**Authors:** Hannah W. Miller, Tammie S.Y. Tam, Katherine S. Ralston

**Affiliations:** Department of Microbiology and Molecular Genetics, University of California, Davis, USA

## Abstract

*Entamoeba histolytica* is the cause of amoebiasis. The trophozoite (amoeba) form of this parasite is capable of invading the intestine, and can disseminate through the bloodstream to other organs. The mechanisms that allow amoebae to evade complement deposition during dissemination have not been well characterized. We previously discovered a novel complement-evasion mechanism employed by *E. histolytica*. *E. histolytica* ingests small bites of living human cells in a process termed trogocytosis. We demonstrated that amoebae were protected from lysis by human serum following trogocytosis of human cells, and that amoebae acquired and displayed human membrane proteins from the cells they ingested. Here, we aimed to define how amoebae are protected from complement lysis after performing trogocytosis. We found that amoebae were protected from complement lysis after ingestion of both human Jurkat T cells and red blood cells, and that the level of protection correlated with the amount of material ingested. Trogocytosis of human cells led to a reduction in deposition of C3b on the surface of amoebae. We asked whether display of human complement regulators is involved in amoebic protection, and found that CD59 was displayed by amoebae after trogocytosis. Deletion of a single complement regulatory protein, CD59 or CD46, from Jurkat cells was not sufficient to alter amoebic protection from lysis, suggesting that multiple, redundant complement regulators mediate amoebic protection. However, exogeneous expression of CD46 or CD55 in amoebae was sufficient to confer protection from lysis. These studies shed light on a novel strategy for immune evasion by a pathogen.

**IMPORTANCE:** *Entamoeba histolytica* is the cause of amoebiasis, a diarrheal disease of global importance. While infection is often asymptomatic, the trophozoite (amoeba) form of this parasite is capable of invading and ulcerating the intestine, and can disseminate through the bloodstream to other organs. Understanding how *E. histolytica* evades the complement system during dissemination is of great interest. Here we demonstrate for the first time that amoebae that have performed trogocytosis (nibbling of human cells) resist deposition of the complement protein C3b. Amoebae that have performed trogocytosis display the complement regulatory protein CD59. Overall, our studies suggest that acquisition and display of multiple, redundant complement regulators is involved in amoebic protection from complement lysis. These findings shed light on a novel strategy for immune evasion by a pathogen. Since other parasites use trogocytosis for cell killing, our findings may apply to the pathogenesis of other infections.

## INTRODUCTION

Amoebiasis remains a disease of global importance. The 2015 Global Burden of Disease Study estimated that it was responsible for 67,900 deaths worldwide that year (1). Its causative agent, *Entamoeba histolytica,* is prevalent in countries with poor sanitation and is spread through feces-contaminated food and water (2). In the rural area of Durango, Mexico, the seroprevalence of *E. histolytica* was found to be as high as 42% (3), and a longitudinal study of children living in an urban community of Dhaka, Bangladesh found that 80% were infected with the parasite by two years of age (4).

While infection with *E. histolytica* is often asymptomatic, it can result in diarrheal disease, colitis and extraintestinal abscesses (5). Following ingestion of the cyst form of the parasite, excystation occurs and the trophozoite stage (amoeba) colonizes the large intestine (6). Amoebae are capable of invading and ulcerating the intestine, causing tissue damage and bloody diarrhea. They can also disseminate through the bloodstream to other organs, most commonly the liver, where they form abscesses that are fatal if left untreated (5). The mechanisms that allow amoebae to evade complement deposition during dissemination have not been well characterized. Pathogenic strains of *E. histolytica* isolated from patients have been shown to be more resistant to complement lysis than nonpathogenic strains (7). It has also been found that the pathogenic amoeba species *E. histolytica* appears to be more resistant to complement than its nonpathogenic relative *E. dispar* (8). *E. histolytica* cysteine proteases can cleave complement components (9–11), and the Gal/GalNAc lectin has been described as a CD59 mimicry molecule (12). However, these mechanisms alone are not sufficient to fully protect amoebae from complement lysis, as trophozoites are readily lysed by human serum *in vitro* (13).

We previously discovered a novel complement-evasion mechanism employed by *E. histolytica* (13). Trogocytosis, or “cell nibbling,” is present in many eukaryotes and occurs in a variety of contexts (14). In *E. histolytica*, trogocytosis is a process in which amoebae ingest small bites of living human cells (15). Our previous work has defined amoebic trogocytosis as both a mechanism for cell killing (15), and more recently, for immune evasion (13). We demonstrated that amoebae were protected from lysis by human serum following trogocytosis of human cells (13), and that amoebae acquired and displayed human membrane proteins from the cells they ingested (13). This work suggested a model in which amoebae incorporate proteins from human cells they eat on their surface, and these proteins in turn inhibit complement lysis.

In the present study, we aimed to define the mechanism by which amoebae are protected from complement lysis after performing trogocytosis. We found that trogocytosis of human cells reduced deposition of the complement protein C3b the surface of amoebae, and that amoebae were protected from complement lysis after ingestion of both human Jurkat T cells and primary red blood cells. We identified the human complement regulatory protein CD59 (protectin) as one of the proteins that is taken from ingested human cells and displayed on the amoebic surface. Deletion of a single complement regulatory protein, CD59 or CD46, from human Jurkat cells was not sufficient to alter conferred protection on amoebae. However, exogeneous expression of a single complement regulator, i.e., CD46 or CD55, in amoebae was sufficient to confer protection from lysis. Overall, these studies suggest that multiple, redundant complement regulators are involved in amoebic protection.

## RESULTS

### Ingestion of beads does not confer protection from complement lysis

We previously showed that protection from complement lysis occurred in amoebae that had performed trogocytosis on living human cells, and that known inhibitors of trogocytosis also inhibited protection from complement lysis (13). Amoebae that had performed phagocytosis of dead human cells did not become protected from complement. Likewise, amoebae that were capable of trogocytosis, but defective in phagocytosis, nonetheless became protected from complement (13). To extend these findings, and to examine the role of ingestion on protection in the absence of host proteins, we allowed amoebae to ingest increasing concentrations of latex beads. Amoebae that ingested beads were no more protected from lysis than control amoebae that did not perform ingestion **(Fig. 1, S1)**. Thus, trogocytosis confers protection from complement lysis, but phagocytosis of dead human cells or latex beads does not confer protection.

**Figure 1:**
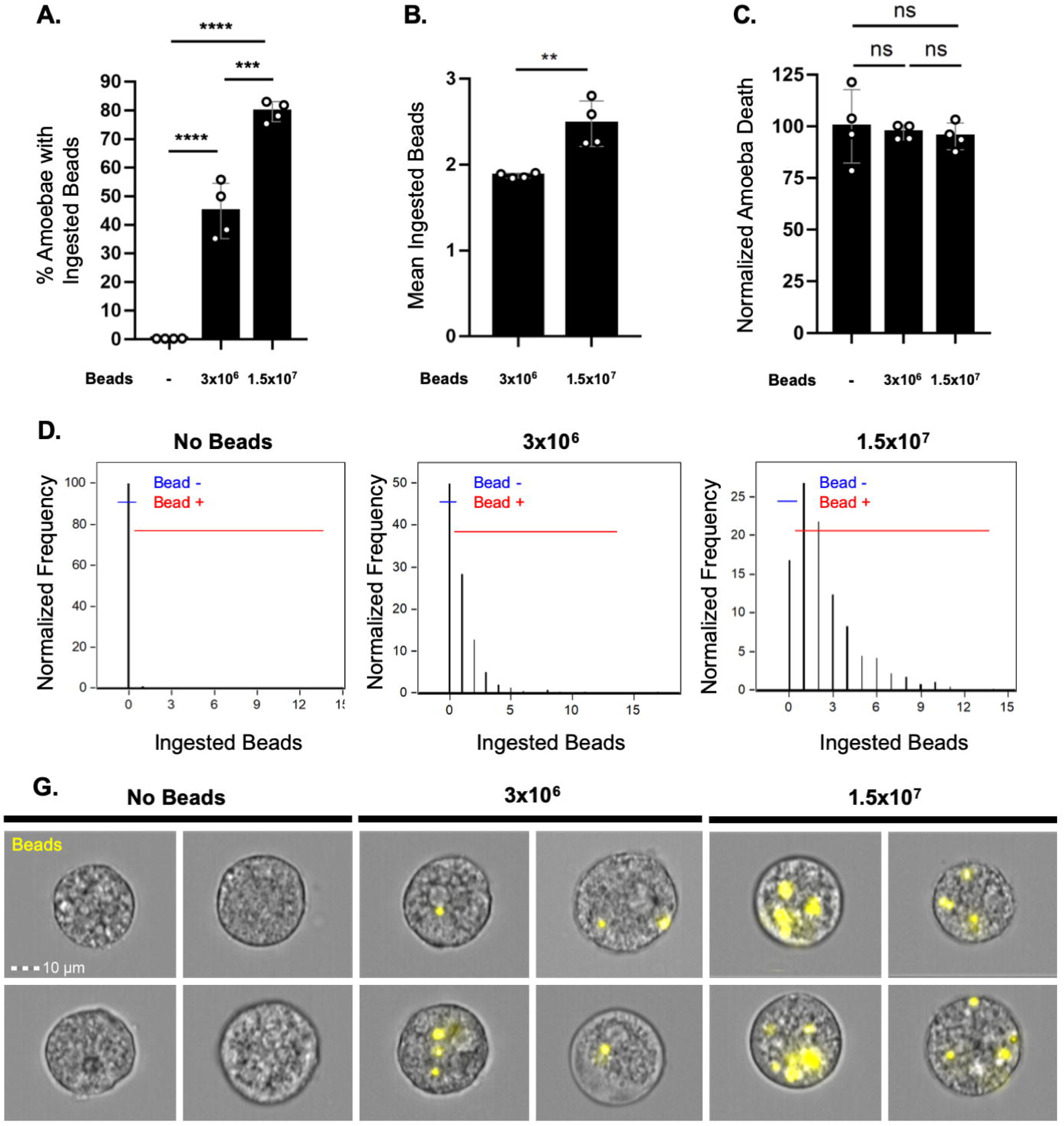
Ingestion of latex beads does not protect amoebae from complement lysis. Amoebae were incubated with 3×10^6^ or 1.5×10^7^ fluorescent latex beads for 1 hour or were incubated without latex beads, and then exposed to human serum. Amoebic viability was determined with Zombie Violet viability dye. Imaging flow cytometry was used to quantify bead ingestion and amoebic viability. **(A)** The percentage of amoebae that had ingested any number of latex beads. **(B)** Mean number of ingested beads among amoebae that had ingested beads. **(C)** Normalized amoeba death following exposure to human serum. Values are normalized to the amoebae that were incubated without latex beads. **(D)** Quantification of the number of ingested beads per amoeba; shown are the data plots from one replicate per sample type. Amoebae that did not perform bead ingestion fall into the “Bead –“ gate, while amoebae that ingested beads fall into the “Bead +” gate. **(G)** Representative images of amoebae that were incubated without beads, or in the presence of 3×10^6^ or 1.5×10^7^ beads. Ingested beads are shown in yellow. Data are from four replicates across two independent experiments.

### Amoebic protection from complement following trogocytosis is dose-dependent

To determine if the amount of trogocytosed human cell material influenced protection, or if any level of trogocytosis was protective, we allowed amoebae to ingest incrementally larger amounts. Amoebae became increasingly protected from complement lysis **(Fig. 2b-c, S2a)** after ingesting higher quantities of live Jurkat cell material **(Fig. 2a, 2g)**. Thus, acquired protection from complement lysis correlates with the amount of human cell material ingested through trogocytosis.

**Figure 2:**
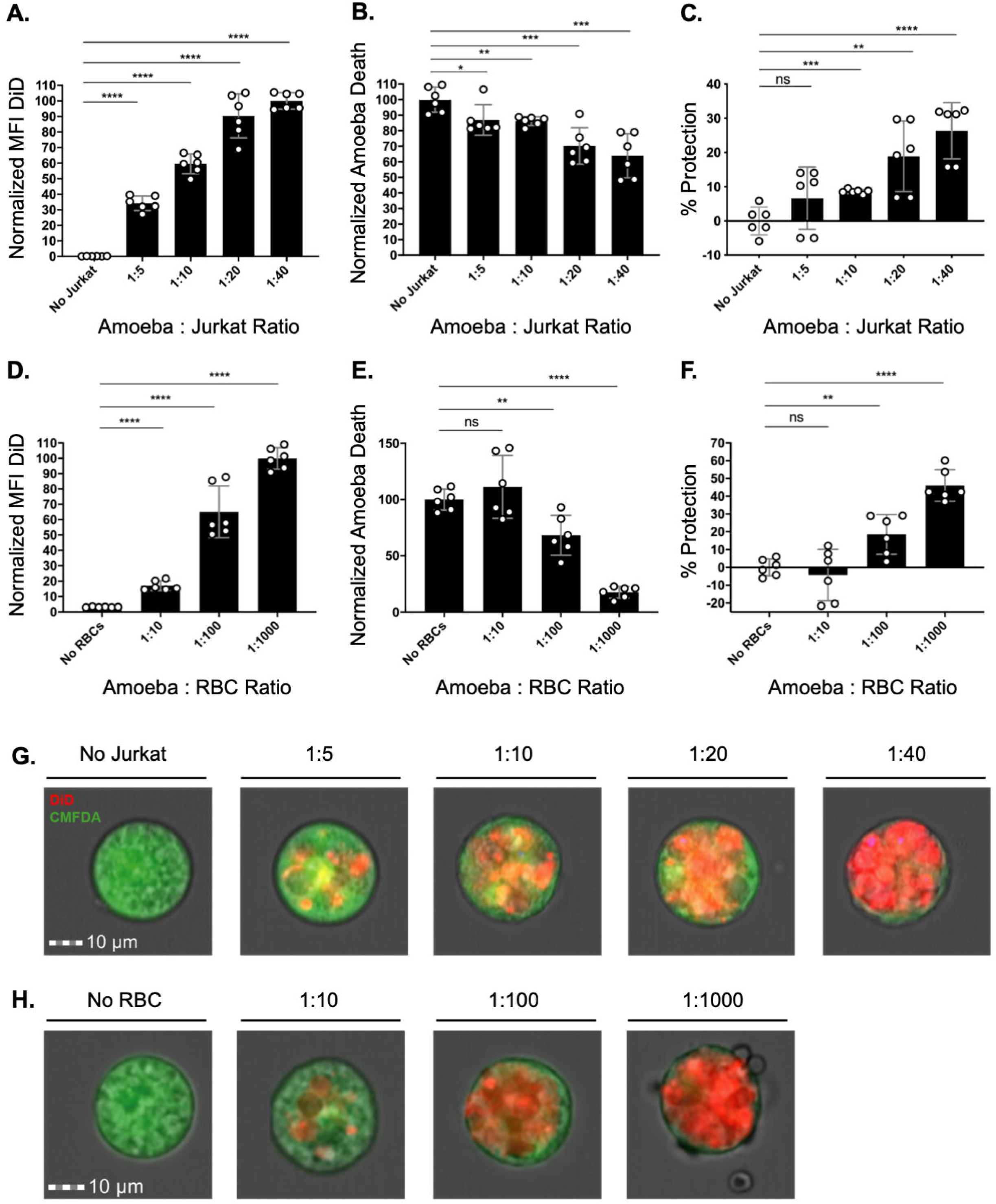
Amoebic protection from complement following trogocytosis is dose-dependent. Amoebae were labeled with CMFDA cytoplasm dye and incubated in the absence of human cells, or with increasing concentrations of human Jurkat cells or primary human red blood cells. Human cells were labeled with DiD membrane dye. Following exposure to human serum, amoeba death was assessed with Zombie Violet viability dye and ingested human cell material was determined by quantifying mean fluorescence intensity (MFI) of DiD present on amoebae. **(A)** Normalized MFI of DiD on amoebae incubated in the absence of Jurkat cells or with increasing concentrations of Jurkat cells. **(B)** Normalized death of amoebae from conditions in panel A. **(C)** Death of amoebae from conditions in panel A, expressed as percent protection. Percent protection was calculated by subtracting the total death of amoebae incubated with human cells from the total death of amoebae incubated in the absence of Jurkat cells. **(D)** Normalized MFI of DiD on amoebae incubated in the absence of red blood cells or with increasing concentrations of red blood cells. **(E)** Normalized death of amoebae from conditions in panel D. **(F)** Death of amoebae from conditions in panel D, expressed as percent protection. **(G)** Representative images of amoebae incubated with increasing concentrations of Jurkat cells. Amoebae are shown in green and ingested human cell material is shown in red. Data were analyzed by imaging flow cytometry and are from 6 replicates across 3 independent experiments. **(H)** Representative images of amoebae incubated with red blood cells. Data were analyzed by imaging flow cytometry and are from 6 replicates across 3 independent experiments.

We next asked if protection could be conferred through trogocytosis of primary human cells. During invasive infections, amoebae breach the intestinal wall and disseminate via the bloodstream. Detection of amoebae containing ingested red blood cells in the stool has previously been used as a diagnostic for invasive disease (16). Therefore, we asked if ingestion of human red blood cells would lead to protection. Amoebae were allowed to ingest increasing numbers of human red blood cells. Increased trogocytosis of red blood cells led to increased protection from complement lysis **(Fig. 2d-f, 2h, S2b)**. These results support a model where the level of protection from complement lysis is proportional to the amount of human cell material that was ingested during trogocytosis.

### Amoebic trogocytosis of human cells inhibits deposition of complement C3b

It can be inferred that lysis of amoebae by human serum is due to complement activity because heat-inactivated serum does not lyse amoebae (13). Here, we formally tested if trogocytosis of human cells led to reduced deposition of complement on the amoebae surface. Amoebae that had been co-incubated with live human cells had less death and less deposited human C3b on their surface than amoebae that were incubated in the absence of Jurkat cells **(Fig. 3a-b, S3)**. Furthermore, live amoebae had less deposited C3b than dead amoebae **(Fig. 3c, 3e)**. Amoebae that had been co-incubated with human cells had less deposited C3b, compared to amoebae that were incubated in the absence of Jurkat cells **(Fig. 3d)**. Therefore, trogocytosis of human cells prevents complement lysis of amoebae and inhibits deposition of complement C3b.

**Figure 3:**
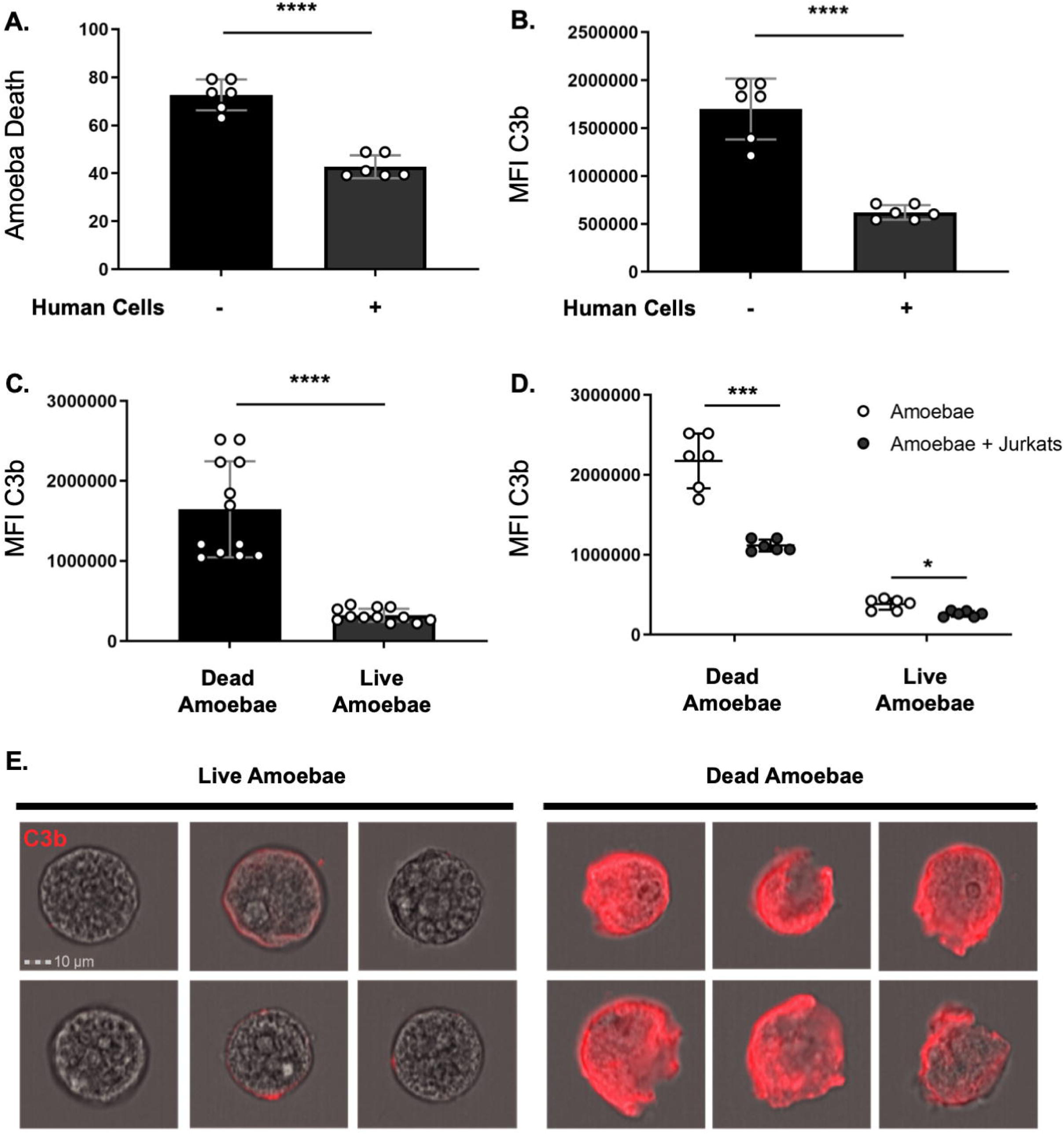
Amoebic trogocytosis of human cells inhibits deposition of complement C3b. Amoebae were incubated in the absence of Jurkat T cells or in the presence of Jurkat cells, and subsequently exposed to human serum. Viability was assessed with Zombie Violet dye. The presence of C3b was detected using a mouse monoclonal antibody to C3b and iC3b. **(A)** Death of amoebae that were incubated in the absence of Jurkat cells or in the presence of Jurkat cells. **(B)** Mean fluorescence intensity of deposited C3b on amoebae. **(C)** Deposited C3b on dead or live amoebae. **(D)** Deposited C3b on dead or live amoebae that had been incubated in the absence of Jurkat cells (open circles) or in the presence of Jurkat cells (filled circles). **(E)** Representative images of C3b deposition (red) on live or dead amoebae. Data were analyzed by imaging flow cytometry and are from 6 replicates across 3 independent experiments.

### Amoebae acquire the complement regulatory proteins CD59 and CD46 from human cells

We hypothesized that protection from complement lysis was due to display of human complement regulatory proteins. We first chose to look at acquisition of the complement regulatory protein CD59 (protectin), a membrane protein that is expressed by both Jurkat cells and human red blood cells (17–19). CD59 inhibits terminal components of the complement cascade and formation of the membrane attack complex (17, 20–22). After amoebae had performed trogocytosis, patches of CD59 were detected on the amoeba surface within five minutes **(Fig. 4a)** and a larger quantity of CD59 patches were detected on the amoeba surface after one hour of trogocytosis **(Fig. 4a-c)**. The patchy/punctate localization pattern was similar to the localization pattern of other human proteins displayed by amoebae after trogocytosis (13). Importantly, the apparent display of human membrane proteins by amoebae was not an artifact of paraformaldehyde fixation, since the appearance of human membrane proteins displayed by amoebae was indistinguishable, whether staining was performed before or after fixation (**Fig. S4** and (13)). Since the heavy chain of the amoeba surface Gal/GalNAc lectin has been implicated as a CD59 mimicry molecule (12, 23), we asked if the CD59 antibody used in these assays cross-reacted with the Gal/GalNAc lectin. CD59 was not detected on control amoebae that had not performed trogocytosis **(Fig. 4a**), showing that the CD59 antibody did not cross-react with the Gal/GalNAc lectin.

**Figure 4:**
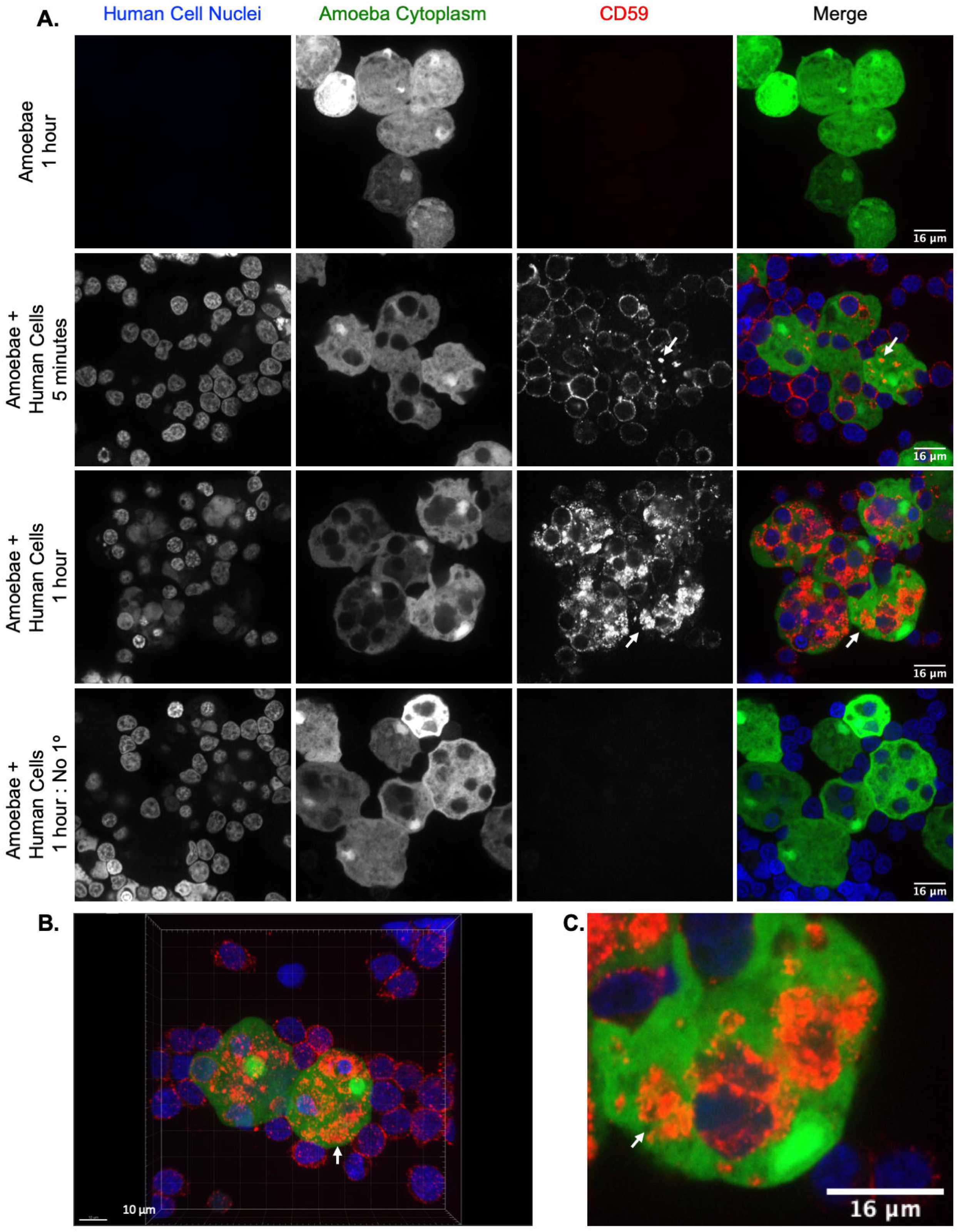
Amoebae acquire and display the complement regulatory protein CD59 from human cells. Amoebae were allowed to perform trogocytosis on human Jurkat T cells for 5 minutes or 1 hour, or were incubated in the absence of Jurkat cells. Human CD59 (red) was detected on the amoebae surface by monoclonal antibody staining. Amoebae were labeled with CMFDA (green) and human cell nuclei were labeled with Hoechst (blue). **(A)** Representative images from amoebae incubated in the absence of Jurkat cells or amoebae that performed trogocytosis on human Jurkat T cells for 5 minutes or 1 hour. Arrows indicate patches of displayed CD59 on the amoeba surface. **(B)** 3D rendering of Z stack images taken from amoebae that were incubated with human Jurkat T cells for 1 hour. **(C)** Zoomed in image of amoebae that were incubated with human Jurkat T cells for 1 hour. Data were analyzed by confocal microscopy. 136 Images were collected from 1 independent experiment.

Imaging flow cytometry analysis was used to quantify amoebic acquisition of CD59. Human cell nuclei are not ingested during trogocytosis (15). Therefore, labeling of human cell nuclei was used to differentiate intact, extracellular human cells from patches of human proteins displayed on amoebae. Patches of displayed human CD59 were detected on amoebae that had performed trogocytosis, but not on amoebae that were incubated in the absence of Jurkat cells **(Fig. 5a-c, S5)**. Amoebae had ∼38% more displayed CD59 after one hour of trogocytosis than after five minutes **(Fig. 5b)**. These findings indicate that amoebae acquired and displayed CD59 within five minutes of trogocytosis, and the quantity of displayed CD59 increased over time.

**Figure 5:**
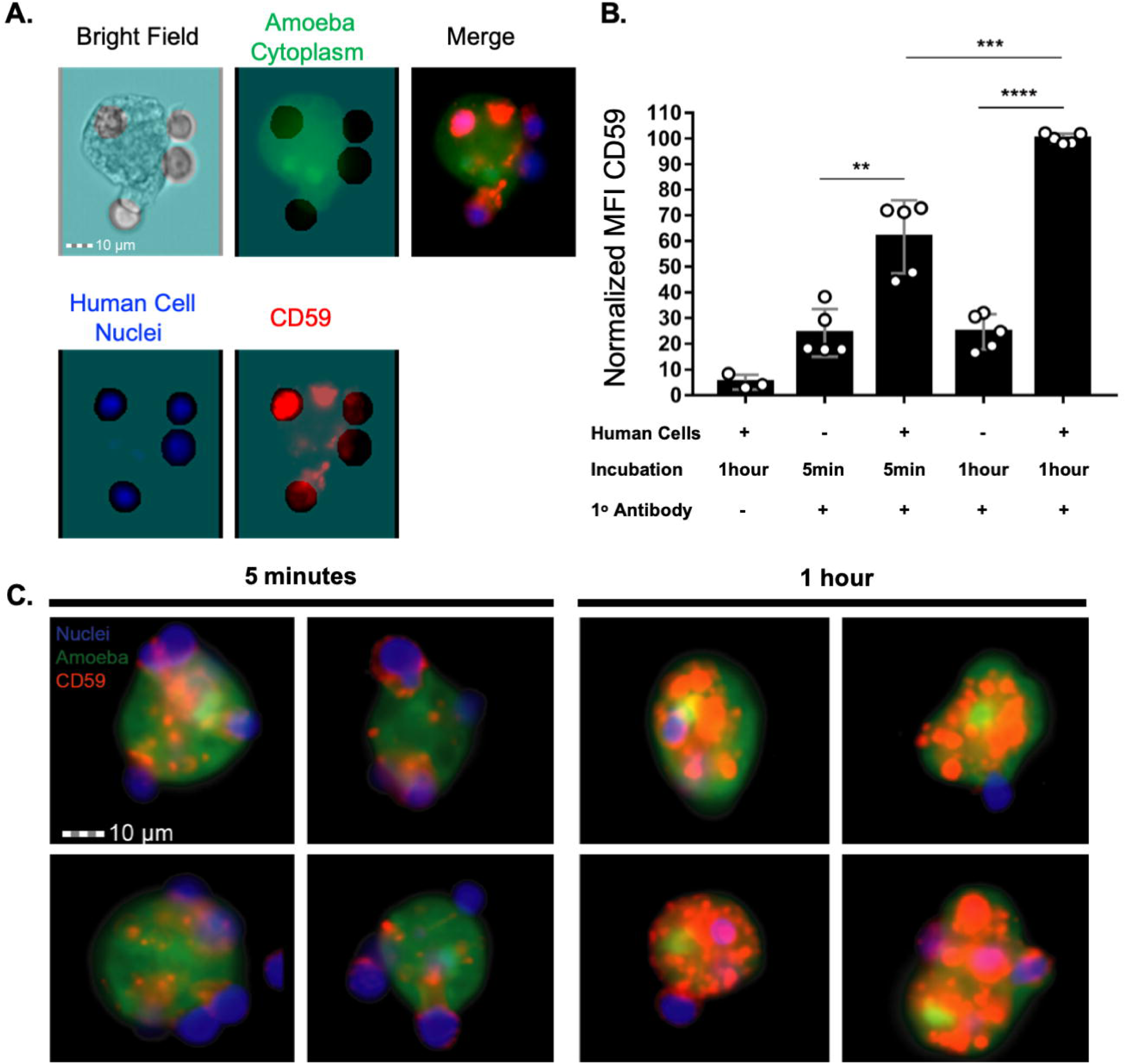
The amount of displayed CD59 increases with increased trogocytosis of human cells. Acquired CD59 molecules were quantified on amoebae that were allowed to perform trogocytosis on human Jurkat T cells for 5 minutes or 1 hour, or were incubated in the absence of Jurkat cells. **(A)** Masking strategy for analysis of displayed CD59 on the amoeba surface. A mask was created in order to allow for the detection of CD59 that overlapped with amoebae, while excluding CD59 on intact human cells attached to amoebae. The mask is displayed in turquoise, as an overlay on the individual images. Amoebae were labeled with CMFDA (green), human cell nuclei were labeled with Hoechst (blue), and CD59 was detected with a monoclonal antibody (red). Extracellular human cell nuclei fluorescence was removed from the masked analysis area. The excluded area around human cell nuclei was then dilated by 4 pixels to include the entire diameter of the intact extra-cellular human cells and associated CD59. CD59 was analyzed in the remaining masked analysis area of each image to allow for analysis of displayed patches of acquired CD59 on amoebae. **(B)** The normalized mean fluorescence intensity of CD59 on amoebae after 5 minutes or 1 hour of trogocytosis or amoebae that were incubated in the absence of Jurkat cells. To normalize the data, samples were normalized to the 1 hour of trogocytosis condition. **(C)** Representative images of amoebae that had performed trogocytosis on human Jurkat T cells for 5 minutes or 1 hour. Arrows indicate displayed CD59. Data were analyzed by imaging flow cytometry and are from 5 replicates across 3 independent experiments. The no primary control condition was performed in 2 of 3 independent experiments and data are from 3 replicates.

To determine if more than one complement regulatory protein was displayed by amoebae after trogocytosis, immunofluorescence was used to detect human CD46. CD46 (membrane cofactor protein) is expressed by human Jurkat cells (24) and acts as a co-factor for serum factor I, which cleaves and inactivates complement components C3b and C4b (25–27). After amoebae had performed trogocytosis, patches of CD46 were detected on the amoeba surface within five minutes **(Fig. S6).** Thus, amoebae acquire and display multiple complement regulatory proteins, after performing trogocytosis of human cells.

### Removal of GPI-anchored surface proteins trends towards restoring complement lysis of amoebae

We next sought to test the effect of removing complement regulatory proteins on conferred protection. CD59 is a glycosylphosphatidylinositol (GPI) anchored protein that can be cleaved with the enzyme phosphatidylinositol-specific phospholipase C (PI-PLC) (17, 18, 20, 22). In addition to removing CD59, treatment with PI-PLC also cleaves the GPI-anchored complement regulatory protein CD55 (decay-accelerating factor) (28, 29). CD55 is expressed by both Jurkat cells and human red blood cells (19, 30) and accelerates the decay of C3 and C5 convertases of the classical and alternative complement pathways (30, 31). Amoebae were allowed to perform trogocytosis or incubated in the absence of Jurkat cells, and then treated with PI-PLC before exposure to human serum. Heat-inactivated PI-PLC was used as a control. PI-PLC treatment appeared to reduce amoebic protection from complement lysis **(Fig. S7, S8a)**. However, this difference was not statistically significant. This is likely to be due to the higher levels of variability in this assay, which were due to the prolonged incubation of amoebae on ice during PI-PLC treatment. Indeed, due to the incubation on ice, the background level of amoeba cell death was much higher than usual. These findings suggested that removal of all GPI-anchored proteins reduces amoebic protection from complement lysis, but did not prove a causal relationship.

### Removal of CD59 and CD46 is not sufficient to sensitize amoebae to complement lysis

Due to the higher levels of variability that we observed with PI-PLC treatment, we next used human cell mutants to test whether individual proteins were required for protection from complement lysis. We tested the requirements for CD59 and CD46. In order to determine if acquisition and display of human CD59 or CD46 molecules was required for protection from complement, we used CRISPR/Cas9 to create human cell knockout mutants that lacked these proteins. Sanger sequencing **(Fig. S9)** and antibody staining of CD59 **(Fig. 6a-b)** and CD46 **(Fig. 6c-d)** showed that knockout mutants were successfully generated. Amoebae that were allowed to perform trogocytosis on control human cells or cells that lacked either CD59 or CD46 were equally protected from complement lysis **(Fig. 6e, S8)**. Additionally, there was no difference in the amount of ingested human cell material **(Fig. 6f)**. These findings reveal that removal of either CD59 or CD46 individually is not sufficient to sensitize amoebae to complement lysis. Furthermore, they hint at redundancy in the mechanism of protection and that amoebae likely acquire and display multiple complement regulatory proteins from the cells they ingest. Display of CD46 or CD55 is sufficient to protect amoebae from complement lysis.

**Figure 6:**
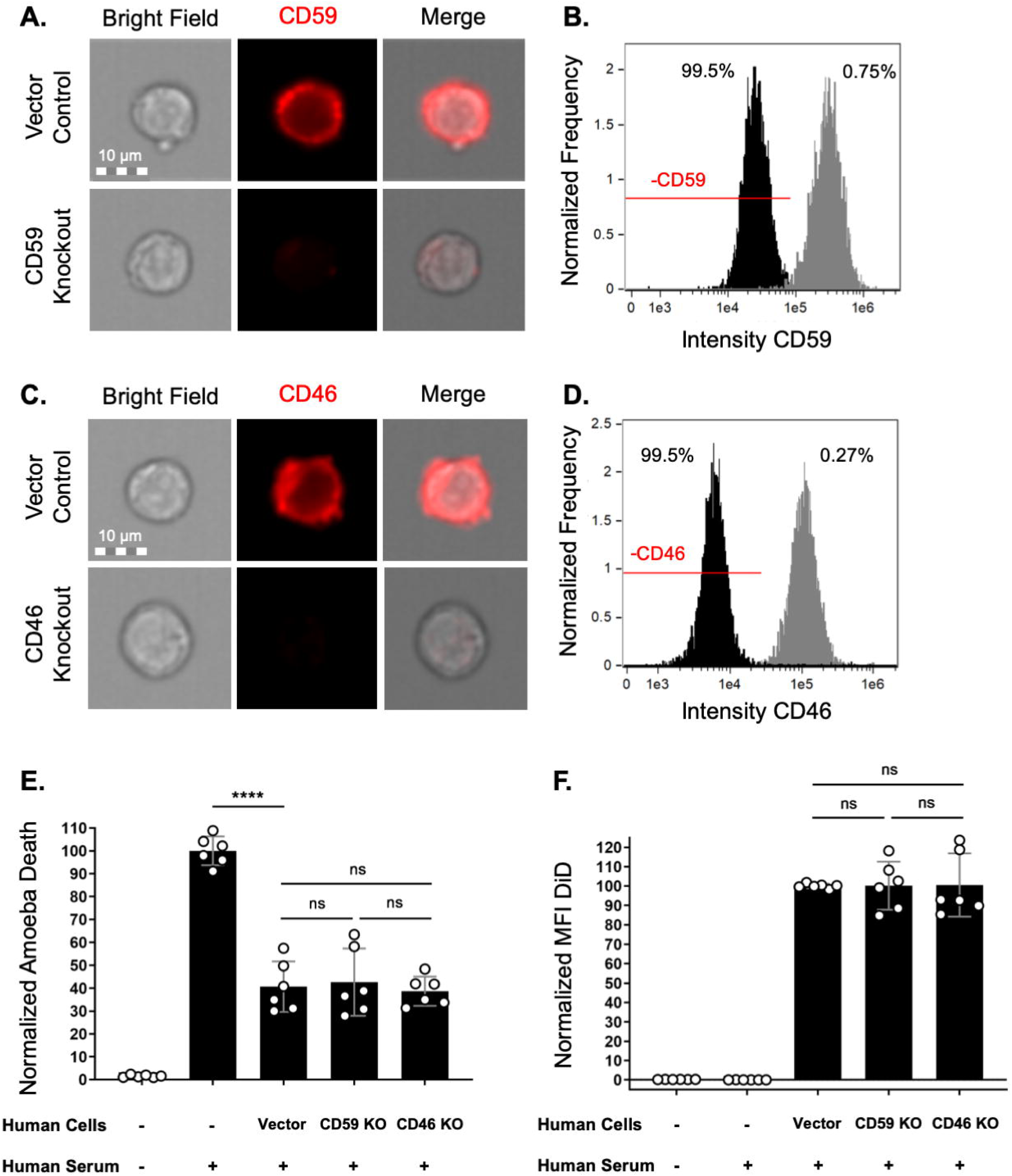
Removal of CD59 and CD46 is not sufficient to sensitize amoebae to complement lysis. **(A-D)** Human Jurkat T cells deficient in CD59 or CD46 were constructed using CRISPR/Cas9. Immunofluorescence and imaging flow cytometry were used to quantify CD59 or CD46. **(A)** Representative images of CD59 antibody staining (red) in vector control human cells or CD59 mutants. **(B)** Intensity of CD59 antibody staining in vector control human cells (gray) or CD59 mutants (black). 99.5% of CD59 mutants were in the CD59-negative gate (“-CD59”), while 0.75% of vector control cells were in this gate. **(C)** Representative images of CD46 antibody staining (red) in vector control human cells or CD46 mutants. **(D)** Intensity of CD46 antibody staining in vector control human cells (gray) or CD46 mutants (black). 99.5% of CD46 mutants were in the CD46-negative gate (“-CD46”), while 0.27% of vector control cells were in this gate. **(E-F)** Amoebae were labeled with CMFDA cytoplasm dye and incubated in the absence of Jurkat cells or with Jurkat cells. Human cell lines were either vector control cells, CD59 KO mutants, or CD46 KO mutants. Human cells were labeled with DiD membrane dye. Following exposure to human serum, amoeba death was assessed with Zombie Violet viability dye and ingested human cell material was determined by quantifying mean fluorescence intensity (MFI) of DiD present on amoebae. **(E)** Normalized death of amoebae. **(F)** Normalized mean fluorescence intensity of DiD on amoebae. Data were analyzed by imaging flow cytometry and are from 6 replicates across 3 independent experiments.

To ask if display of human complement regulatory proteins was sufficient to protect amoebae from lysis, we created amoebae that exogenously expressed either human CD46 or CD55 **(Fig. 7, S2a).** Expression of CD46 or CD55 was detectable with RT-PCR analysis **(Fig. 7a).** Immunofluorescence analysis showed that CD46 or CD55 proteins were surface-localized in amoebae, as they were detected with antibody staining of non-permeabilized cells **(Fig. 7b-j, S2a).** Localization of the *E. histolytica* surface Gal/GalNAc lectin was used as a control **(Fig. 7d, 7g, 7j),** and all transfected amoebae had equivalent amounts of Gal/GalNAc lectin. CD46 or CD55 expression was at a relatively low level **(Fig. 7b-c, 7e-f),** just above background fluorescence of vector control amoebae. Additionally, CD46 or CD55 expression was markedly lower than the expression level of the Gal/GalNAc lectin **(Fig. 7d, 7g).** When exposed to human serum, amoebae that expressed human CD46 or CD55 were more protected from lysis than vector control amoebae **(Fig. 8, S8b)**. Therefore, display of a single complement regulatory protein is sufficient to protect amoebae from complement lysis.

**Figure 7:**
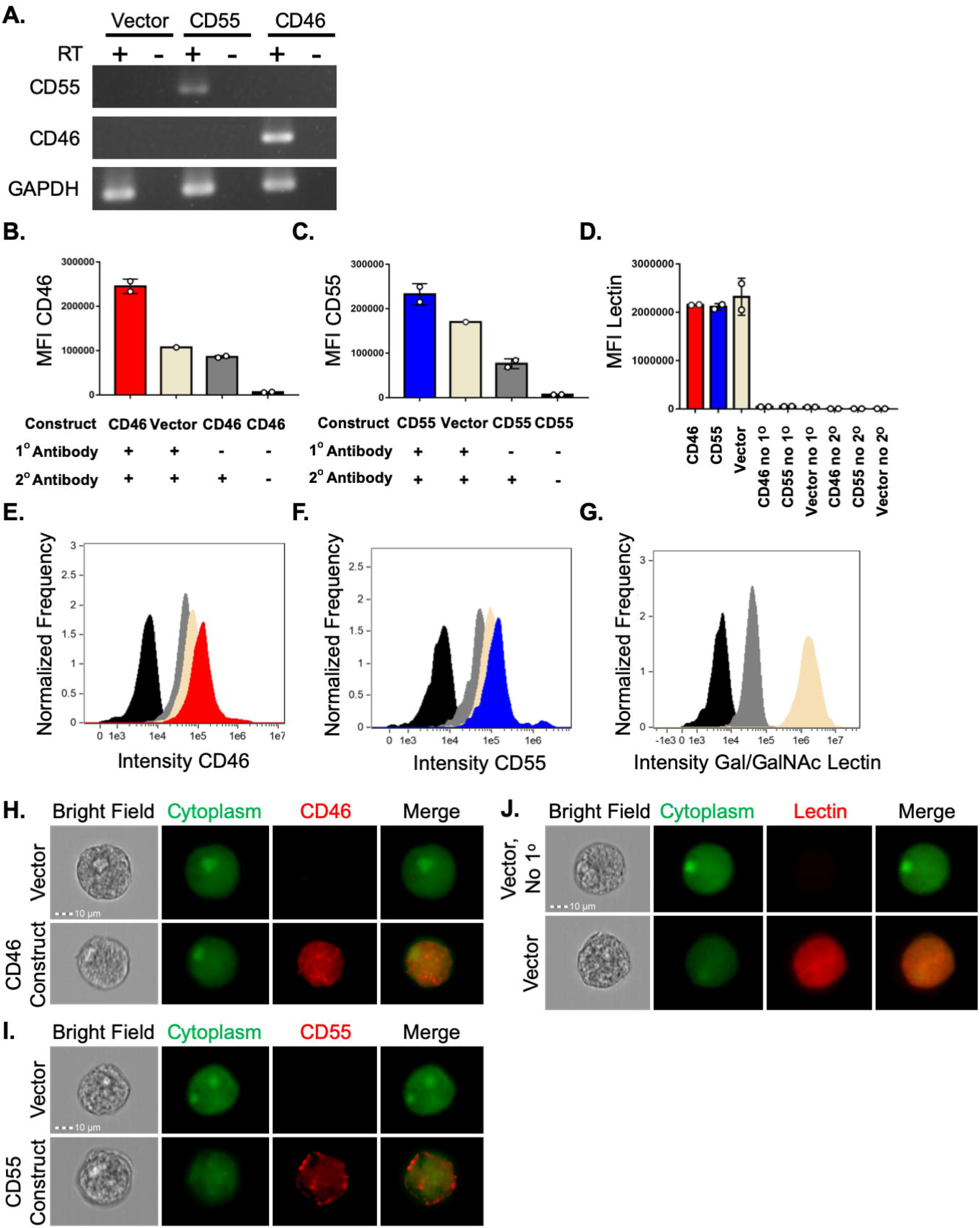
Amoebae transfected with expression plasmids display human CD46 or CD55. Amoebae were transfected with expression constructs for human CD46 or human CD55, or the expression plasmid backbone as a negative control. **(A)** RT-PCR analysis with primers specific for human CD46, human CD55, or amoebic GAPDH. Samples were incubated with or without reverse transcriptase (RT +/-), to control for DNA contamination. **(B-J)** Amoebae were labeled with a cytoplasmic dye (CMFDA) and immunofluorescence was performed without permeabilization. **(B)** Mean fluorescence intensity of human CD46 antibody staining. Amoebae were stably transfected with a plasmid for expression of human CD46 (red) or vector backbone (beige), and were stained using a CD46 primary antibody and a far red secondary antibody. Amoebae expressing human CD46 were also incubated without primary antibody (grey), or without any antibodies (black). **(C)** Mean fluorescence intensity of human CD55 antibody staining. Amoebae were stably transfected with a plasmid for expression of human CD55 (blue) or vector backbone (beige), and were stained using a CD46 primary antibody and a far red secondary antibody. Amoebae expressing human CD55 were also incubated without primary antibody (grey), or without any antibodies (black). **(D)** Mean fluorescence intensity of amoebic Gal/GalNAc lectin antibody staining. Amoebae were stably transfected with a plasmid for expression of human CD46 (red), or human CD55 (blue), or vector backbone (beige), and were stained using a Gal/GalNAc lectin primary antibody and a far red secondary antibody. Amoebae were also incubated without primary antibody, or without any antibodies. **(E-G)** Histograms corresponding to the mean fluorescence intensity data shown in panels B-D. Panel G shows antibody staining of vector control amoebae, stained with both antibodies (beige), without primary antibody (grey), or without any antibodies (black). (H-J) Representative images corresponding to the analysis shown in panels B-G. Data shown in panel A are representative of two replicates. Data shown in panels B-J were analyzed by imaging flow cytometry are from 3-4 replicates across 2 independent experiments; the data from one independent experiment are shown.

**Figure 8:**
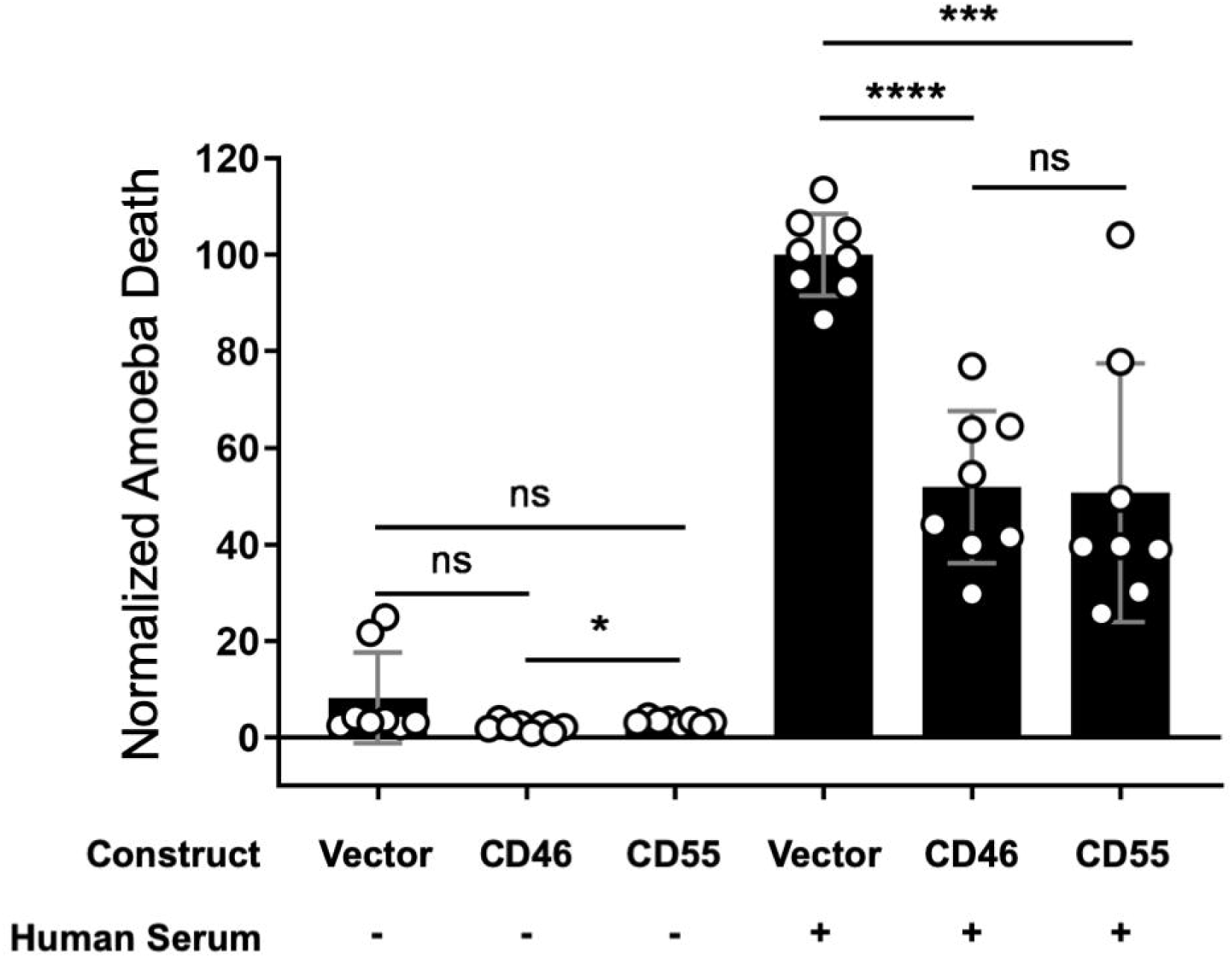
Display of human CD46 or CD55 is sufficient to protect amoebae from complement lysis. Amoebae were stably transfected with CD46 or CD55 expression constructs, or vector backbone. Following exposure to either media or human serum, amoeba viability was assessed using Zombie Violet viability dye and imaging flow cytometry. Amoebic death was normalized to vector control amoebae that were incubated with human serum. These data are from 8 replicates across 4 independent experiments.

## DISCUSSION

Our findings reveal that amoebae are protected from complement lysis through trogocytosis of human cells and that trogocytosis leads to less deposited C3b on the amoeba surface. Amoebae acquire protection from both human Jurkat cells as well as primary red blood cells and conferred protection is proportional to the amount of ingested human cell material. Trogocytosis of human cells results in the display of the complement regulatory protein CD59 on the amoebae surface. Finally, although removal of the individual complement regulatory proteins, CD59 or CD46 was not sufficient to sensitize amoebae to complement lysis, the display of an individual complement regulatory protein, CD46 or CD55, was sufficient to protect amoebae from complement lysis.

Many studies have shown that cancer cells overexpress complement regulatory molecules, allowing them to evade complement lysis. However, it should be noted that expression levels vary widely between cell types and between individual studies (32). Removal of GPI-anchored proteins, which include the complement regulatory proteins CD59 and CD55 but not CD46, with PI-PLC treatment resulted in enhanced susceptibility of cancer cells to complement lysis (28, 29). This is consistent with our finding that PI-PLC treated amoebae that had undergone trogocytosis of human cells, and been made resistant to complement, became more sensitive to complement lysis. Treatment with PI-PLC removes other GPI-anchored proteins in addition to CD59 and CD55, so we cannot rule out the possibility that loss of other membrane proteins contributed to the loss of complement resistance in our PI-PLC treated amoebae.

Amoebae that were engineered to express complement regulatory proteins became protected from lysis by human complement. We were unable to express human CD59 in amoebae, as although stably transfected amoebae with a CD59 expression construct were successfully recovered, these transfectants grew slowly and could not be maintained over time. While CD59 expression was detectable by RT-PCR, it is possible that the CD59 protein mis-localized in amoebae, or otherwise led to deleterious effects. CD46 and CD55 were each appropriately localized to the amoeba cell surface, as evidenced by immunofluorescence staining, and by the functional protection from complement lysis that they conferred. Notably, CD46 and CD55 were not greatly overexpressed in amoebae. When using immunofluorescence, CD46 and CD55 staining levels were just above the background levels of fluorescence seen in vector control amoebae. Thus, these proteins led to protection from complement lysis without dramatically high levels of expression. Only a single complement regulator at a low expression level was needed for protection from lysis by complement. These findings fit with the generally relatively low endogenous expression levels of complement regulatory proteins in human cells, in that low levels of these proteins are sufficient to inhibit the complement pathway.

There have been several studies that examined the efficacy of using blocking antibodies against complement regulatory proteins to enhance lysis of cancer cells. Results from blocking individual proteins have been highly variable in different cell types but blocking one or more proteins was effective in many cases. Using antibodies against CD46 was rarely effective, however antibodies against CD59 and CD55 often were effective in enhancing complement lysis (32). Additionally, there appears to be an additive effect when multiple proteins are targeted at once. Cervical carcinoma cells were rendered more susceptible to complement after treatment with blocking antibodies to CD59 and CD55, and lysis was increased further when cells were treated with both antibodies (33). Similarly, breast carcinoma cells, were more easily lysed after treatment with anti-CD59 and anti-CD55 antibodies, but a much higher degree of lysis was achieved when a mixture of anti-CD59, anti-CD55, and anti-CD46 antibodies was used (34). Since neither removal of CD46 or CD59 was sufficient to sensitize amoebae to complement lysis, it is possible that amoebic display of multiple different complement regulators enables protection from lysis. In our assays, we used a 1:40 ratio of amoebae to Jurkat cells, a ratio that led to a high level of protection from complement lysis. It is possible that using fewer human cells, to lead to a suboptimal level of protection from complement lysis, might have more sensitively revealed differences in amoebic protection from complement lysis due to subtle perturbations like the loss of individual human complement regulators. While removal of CD46 or CD59 did not sensitize amoebae to complement lysis, these proteins could still contribute to amoebic complement protection, as part of a collection of multiple redundant, complement regulators that protect amoebae from lysis. Indeed, experiments in which amoebae were engineered to express complement regulators showed that display of CD46 or CD55 was sufficient to protect amoebae from lysis, supporting the relevance of these proteins in protecting amoebae that have performed trogocytosis.

Our results support a model whereby amoebae are protected from serum lysis in the blood through trogocytosis of red blood cells they encounter there, and potentially from other cells they encounter before reaching the bloodstream. Amoebae likely acquire and display multiple complement regulatory proteins from the human cells that they ingest, which then leads to less C3b deposition, and protection from complement lysis.

## MATERIALS AND METHODS

### Cell culture

*E. histolytica* trophozoites (HM1:IMSS) from ATCC (amoebae) were cultured as described previously (13, 35). Amoebae were maintained in glass tissue culture tubes at 35°C in TYI-S-33 medium supplemented with 15% heat-inactivated adult bovine serum (Gemini Bio-Products), 2.3% Diamond vitamin Tween 80 solution (40×; Sigma-Aldrich), and 80 U/ml penicillin, 80 µg/ml streptomycin (Gibco). Amoebae were expanded in T25 un-vented tissue culture flasks and harvested when flasks reached 80% confluence. When used in serum lysis or immunofluorescence assays, amoebae were resuspended in M199s medium (Gibco medium M199 with Earle’s salts, L-glutamine, and 2.2 g/liter sodium bicarbonate, without phenol red) supplemented with 0.5% bovine serum albumin (BSA), 25 mM HEPES, and 5.7 mM L-cysteine.

Human Jurkat T cells (clone E6-1) from ATCC were grown in vented T25 tissue culture flasks at 37°C and 5% CO2 as previously described (13). Jurkat cells were cultured in RPMI 1640 medium (Gibco; RPMI 1640 with L-glutamine and without phenol red) supplemented with 10% heat-inactivated fetal bovine serum (Gibco), 100 U/ml penicillin, 100 µg/ml streptomycin, and 10 mM HEPES. Jurkat cells were expanded in T75 vented tissue culture flasks and harvested when cell density reached between 5 × 10^5^ and 2 × 10^6^ cells/ml. Jurkat cells were resuspended in M199s medium for use in serum lysis or immunofluorescence assays.

Single donor human red blood cells separated from whole blood by centrifugation and negative for the presence of human immunodeficiency virus-1 (HIV-1), HIV-2, HIV-1 antigen or HIV-1 nucleic acid test, hepatitis B surface antigen, hepatitis C virus, syphilis, and alanine aminotransferase test were purchased from Innovative Research (Cat# IWB3CPDA1UNIT). Red blood cells were stored at 4°C and resuspended in M199s medium prior to use in serum lysis assays.

### DNA constructs

For creation of Jurkat cell CRISPR knockout mutants, guide RNAs (gRNA) to human CD59 or CD46 were cloned into a pX330-U6-Chimeric_BB-CBh-hSpCas9 plasmid backbone (pX330-U6-Chimeric_BB-CBh-hSpCas9 was a gift from Feng Zhang (Addgene plasmid # 42230; http://n2t.net/addgene:42230 ; RRID:Addgene_42230) (36)). Guide RNAs were created by using the sequences designed by Thielen et. al, 2018 (37). The gRNA oligos used are presented in Table 1 with BbsI restriction enzyme overhangs shown in bold. Guide RNAs were cloned into the pX330-U6-Chimeric_BB-CBh-hSpCas9 plasmid backbone using a modified version of the Zhang Lab General Cloning Protocol (Addgene http://www.addgene.org/crispr/zhang/). Briefly, the px330 plasmid backbone was digested with BbsI restriction enzyme (FastDigest BpiI: ThermoFisher Scientific). Guide RNA oligos containing BbsI overhangs were then phosphorylated and annealed. Next, annealed oligos were ligated into the pX330 plasmid backbone and NEB 5-alpha Competent *E. coli* were transformed (New England Biolabs). Positive colonies were screened by restriction digest. Plasmids with the correct inserts were confirmed by Sanger sequencing using the “U6” Universal Primer from GENEWIZ (LKO.1 5’) which is located in the human U6 promoter **(Table 1)**.

**Table 1:**
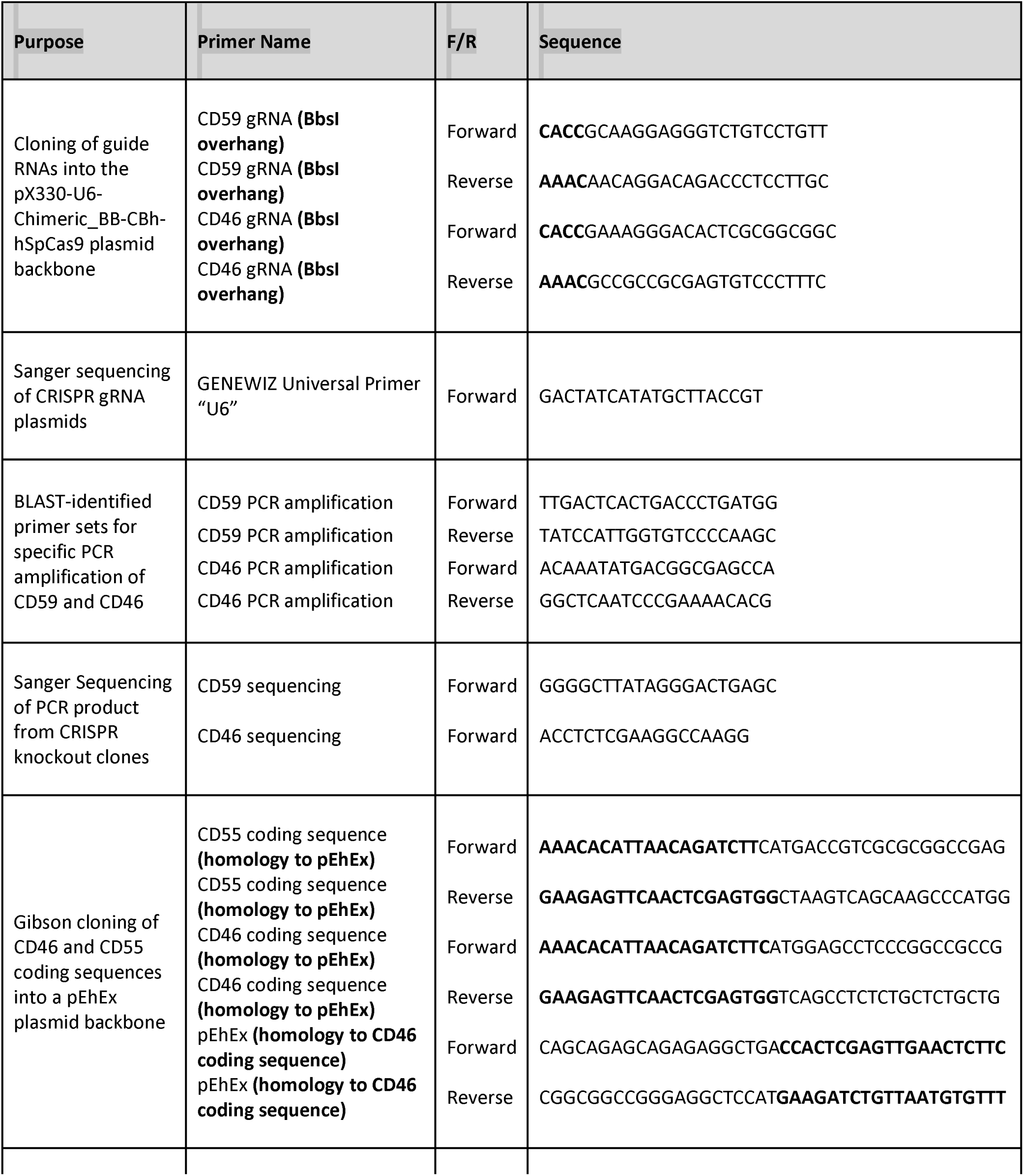
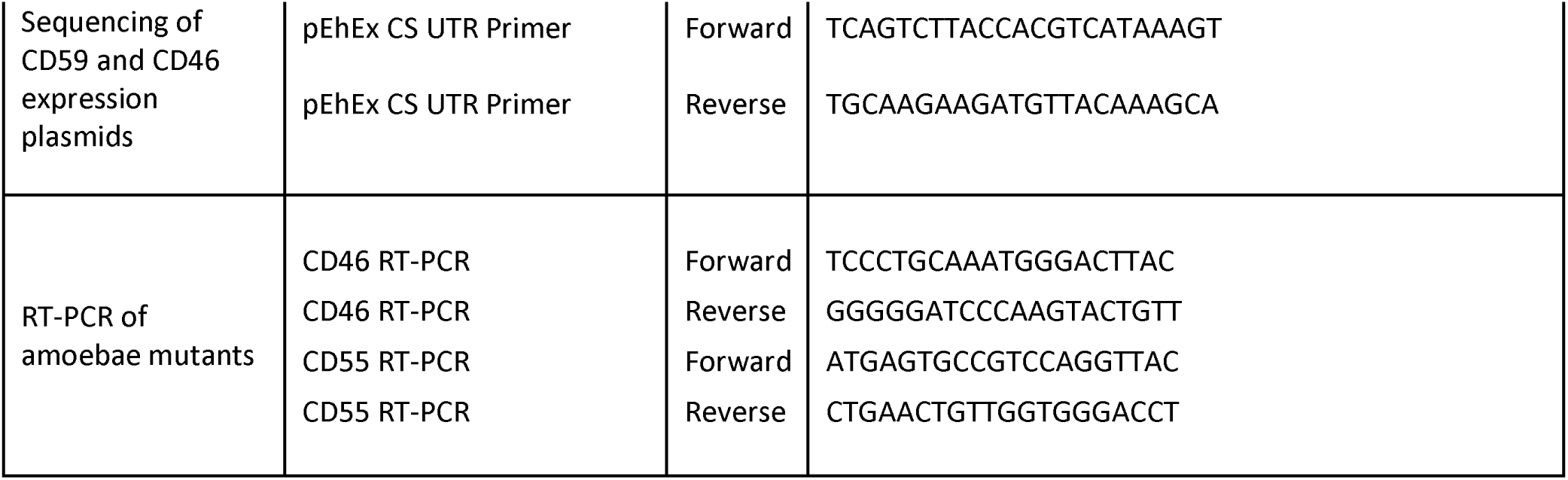
Primers used in these studies.

To generate amoebic expression constructs for exogeneous expression of human proteins, the coding regions of human CD46 or CD55 were amplified from human Jurkat cells. Briefly, RNA was extract from Jurat cells using the Direct-zol RNA MiniPrep Plus kit (Zymo Research) and cDNA was prepared. Next, the coding sequences for human CD46 or CD55 were amplified from Jurkat cDNA by PCR using primers described in **Table 1.** The amplified CD46 or CD55 inserts were then cloned into the *E. histolytica* expression plasmid pEhEx backbone (38) using the Gibson Assembly Ultra Kit (VWR). Primers used for Gibson cloning are described in **Table 1.** For creation of the CD46 expression plasmid, both the pEhEx backbone and the CD46 insert were PCR amplified with primers that added 20 base pairs corresponding to the ends of the insert or backbone, respectively. For creation of the CD55 expression plasmid, the pEhEx backbone was digested using XhoI (New England Biolabs) and the CD55 insert was PCR amplified with primers that added 20 base pairs corresponding to the ends of the backbone. NEB 5-alpha Competent *E. coli* were transformed, and positive colonies were screened by restriction digest analysis. Sanger sequencing was used to assess the sequences of the inserts and flanking vector sequences, using primers located in the Cysteine synthase A gene [EHI_024230] 5’ and 3’ UTR regions **(Table 1)**. The CD46 gene insert corresponds to the mRNA transcript variant d (GenBank accession number: NM_153826) and the CD55 gene insert corresponds to mRNA transcript variant 1 (GenBank accession number: NM_000574).

### Jurkat T cell CRISPR/Cas9 mutants

Human Jurkat T cells were transfected with pX330 plasmids containing the CD59 gRNA, the CD46 gRNA, or the pX330 backbone as a control. Plasmid DNA was isolated in an endotoxin-free manner (GenElute™ HP Endotoxin-Free Plasmid Maxiprep Kit; Sigma-Aldrich) and concentrated using paramagnetic beads (HighPrep PCR Clean Up System; Magbio). Jurkat T cells were transfected using the Neon Transfection System (Invitrogen) with the 10 µl tip and 24 well plate format. Cells were prepared, and the Neon Transfection System was used according to the manufacturer’s instructions. The transfection conditions used were as follows: volts = 1050, width = 30 and pulse # = 2.

When creating the CD59 and CD46 knockout mutants, 1.8 µg of plasmid DNA was used with 1 x 10^5^ cells per transfection reaction and two reactions were performed for each plasmid and transferred to 1 well of a 24 well plate. Transfection efficiency was calculated by separately transfecting an enhanced green fluorescent protein (EGFP) expression plasmid pcDNA3-EGFP (pcDNA3-EGFP was a gift from Doug Golenbock (Addgene plasmid # 13031 ; http://n2t.net/addgene:13031 ; RRID:Addgene_13031)) in parallel. Percentage of EGFP expressing cells was calculated after 24 hours by fixing samples with 4% paraformaldehyde (Electron Microscopy Sciences) and analyzing with imaging flow cytometry. To generate CD59 expressing Jurkat cells, 2 µg of plasmid DNA was used with 2 x 10^5^ cells per transfection reaction and two reactions were performed for each plasmid and transferred to 1 well of a 24 well plate. Clonal lines of Jurkat cell mutants were obtained by limiting dilution in 96 well plates.

Clonal lines were screened for knockout by labeling with primary mouse monoclonal antibodies to CD59 (clone MEM-43/5; Abcam) or CD46 (clone C-10; Santa Cruz Biotechnology) and a Cy™5 AffiniPure Goat Anti-Mouse secondary antibody (Jackson ImmunoResearch Laboratories, Inc). Samples were analyzed by imaging flow cytometry. Knockout was confirmed in positive clones by isolating genomic DNA using the Quick-DNA Miniprep kit (Zymo Research) and polymerase chain reaction (PCR) amplifying regions of either CD59 or CD46. Primer sets had been identified in BLAST as specific to these genes **(Table 1)**. Next, purified PCR product was sequenced with primers upstream of the predicted CRISPR/Cas9 cut site **(Table 1)** and knockout was confirmed.

### Characterization of amoebae exogenously expressing human CD46 or CD55

Amoebae were transfected as described previously (13). Briefly, amoebae were transfected with 20 µg of the human CD46 or CD55 expression construct, or the pEhEx plasmid backbone using Attractene transfection reagent (Qiagen). Stable transfectants were initially selected at 3 µg/ml Geneticin (ThermoFisher Scientific), and then maintained at 6 µg/ml Geneticin. Before use in immunofluorescence assays and serum lysis experiments, drug selection was gradually increased from 6 µg/ml to 48 µg/ml Geneticin, to increase plasmid copy number. Amoebae were maintained at 48 µg/ml Geneticin for 24 hours before experiments were performed.

Expression of human CD46 and CD55 was confirmed using RT-PCR analysis of amoebae maintained at 3 µg/ml Geneticin. RNA was extracted using the Direct-zol RNA MiniPrep Plus kit (Zymo Research) and cDNA was made. Primers used for RT-PCR are shown in Table 1. RT-PCR was performed as described (35).

### Imaging flow cytometry

To detect human CD59 displayed on amoebae following trogocytosis, Jurkat cells were with labeled Hoechst 33342 dye (Invitrogen) at 5 µg/ml for 30 minutes at 37°C. Amoebae were washed and resuspended in M199s medium and labeled with CellTracker green 5-chloromethylfluorescein diacetate (CMFDA; Invitrogen) at 186 ng/ml for 10 minutes at 35°C. Amoebae and Jurkat cells were then washed and resuspended in M199s medium. Amoebae were resuspended at 4 x 10^5^ cells/ml and Jurkat cells were resuspended at 1.6 x 10^7^ cells/ ml for a 1:40 amoeba: Jurkat cell ratio. Amoebae and Jurkat cells were co-incubated for either 5 minutes or 1 hour, or amoebae were incubated in the absence of Jurkat cells. Cells were fixed with 4% PFA for 30 minutes at room temperature. Samples were blocked in 1 x PBS containing 0.1% Tween20 (Sigma-Aldrich) (1 x PBST), 5% BSA and 20% Normal Goat Serum (Jackson ImmunoResearch Laboratories, Inc) for 1 hour at room temperature on a rocker. Next, samples were labeled with primary mouse monoclonal antibodies to CD59 (clone MEM-43/5; Abcam) diluted 1:50 in blocking solution at 4°C overnight on a rocker. Samples were washed with 1 x PBST and labeled with a Cy5 AffiniPure Goat Anti-Mouse secondary antibody (Jackson ImmunoResearch Laboratories, Inc) stored in 50% glycerol (Sigma-Aldrich). The secondary antibody was diluted 1:100 in blocking solution, for a final dilution of 1:200, and incubated for 3 hours at room temperature on a rocker. Lastly, samples were washed with 1 x PBST, resuspended in 50 µl 1 x PBS, and run on an Amnis ImageStreamX Mark II. 10,000 events were collected for conditions where amoebae were incubated in the absence of Jurkat cells and 100,000 events were collected for conditions where amoebae were incubated with Jurkat cells. Data are from 5 replicates across 3 independent experiments. The no primary control condition was performed in 2 of 3 independent experiments and data are from 3 replicates. See Figure S5 for the analysis gating scheme.

To detect human CD46 displayed on amoebae following trogocytosis, amoebae and Jurkat cells were labeled in the same manner as for detection of CD59. Amoebae and Jurkat cells were co-incubated for 5 minutes or amoebae were incubated in the absence of Jurkat cells. Samples were labeled with a primary mouse monoclonal antibody to human CD46 (Clone C-10; Santa Cruz Biotechnology) at a 1:10 dilution, followed by a Cy5 AffiniPure Goat Anti-Mouse secondary antibody (Jackson ImmunoResearch Laboratories, Inc). Amoebae were gated on by size and 10,000 events were collected per sample. Data are from 2 replicates and 1 independent experiment.

For detection of CD46, CD55, or Gal/GalNAc lectin expression on amoebae exogenously expressing CD46 or CD55, amoebae were labeled with CMFDA and prepared in the same manner as for detection of displayed human CD59. Following fixation, samples were blocked overnight at 4°C on a rocker. Samples were then labeled with primary mouse monoclonal antibodies to human CD46 (Clone C-10; Santa Cruz Biotechnology) human CD55 (Clone NaM16-4D3; Santa Cruz Biotechnology) or the *E. histolytica* Gal/GalNAc lectin (Clone 7F4; the Gal/GalNAc lectin antibody was a gift from William A. Petri, Jr., University of Virginia (39)) overnight at 4°C on a rocker. The CD46 and CD55 primary antibodies were diluted 1:10 and the Gal/GalNAc lectin primary antibody was diluted 1:50 in blocking solution. Samples were labeled with secondary antibody as described above. Amoebae were gated on by size and 10,000 events were collect per sample. Data are from 3-4 replicates from 2 independent experiments. See Figure S2a for the analysis gating scheme.

### Confocal microscopy

To detect displayed CD59 on the amoeba surface after trogocytosis, cells were labeled for confocal microscopy as described previously (13). Amoebae were prepared in the same manner as the for the detection of displayed CD59 using imaging flow cytometry (above) with an approximate fourfold increased concentration of CMFDA. After labeling with antibodies, samples were mounted using Vectashield (Vector Laboratories) on Superfrost Plus microslides (VWR) with glass coverslips. Slides were imaged on an Intelligent Imaging Innovations hybrid Spinning Disk Confocal-TIRF-Widefield Microscope. 136 Images were collected from 1 independent experiment. FIJI software was used for image analysis (40).

For detection of displayed biotinylated human cell membrane proteins, amoebae were left unstained in some experiments or labeled with CMFDA as described above. Human Jurkat cells were biotinylated as described previously (13). Briefly, Jurkat cells were biotinylated with EZ-Link Sulfo-NHS-SS-Biotin (ThermoFisher Scientific) at 480 µg/ml in 1× PBS for 25 min at 4°C. The reaction was quenched using 100 mM Tris-HCl (pH 8) and washed with washed in 100 mM Tris-HCl (pH 8) before being resuspending M199s medium. Amoebae were incubated with biotinylated Jurkat cells at a 1:5 ratio for 2 - 5 minutes and then immediately placed on ice. Next, samples were stained with Alexa Fluor 633 conjugated streptavidin (Invitrogen) at 20 µg/ml for 1 hour at 4°C, prior to fixation with 4% paraformaldehyde for 30 minutes at room temperature. In some experiments, samples were labeled with DAPI (4, 6-diamidino-2-phenylindole; Sigma-Aldrich) for 10 minutes at room temperature following fixation. Samples were mounted on either poly-lysine (Sigma-Aldrich) or collagen (collagen I, rat tail; Gibco) coated coverslips for 1 hour. Samples were mounted with Vectashield on glass slides, and slides were imaged on an Olympus FV1000 laser point-scanning confocal microscope.

### Serum lysis assays

For experiments where amoebae were incubated with increasing numbers of Jurkat cells, amoebae were washed and resuspended in M199s medium and labeled with CMFDA at 186 ng/ml for 10 minutes at 35°C. Jurkat cells were washed and labeled in M199s with DiD at 21 µg/ml for 5 minutes at 37°C and 10 minutes at 4°C. Amoebae were washed and resuspended at 4 x 10^5^ cells/ml. Jurkat cells were washed a resuspended at 2 x 10^6^ cells/ml, 4 x 10^6^ cells/ml, 8 x 10^6^ cells/ml and 1.6 x 10^7^ cells/ml. Amoebae were incubated in the absence of Jurkat cells or the presence of Jurkat cells at a 1:5, 1:10, 1:20, and 1:40 ratio for 1 hour at 35°C. Next, samples were resuspended in 100% normal human serum (pooled normal human complement serum; Innovative Research Inc.) supplemented with 150 µM CaCl_2_ and 150 µM MgCl_2_ for 30 minutes at 35°C as described previously (13). Following exposure to human serum, samples were resuspended in M199s medium and labeled using Zombie Violet Fixable Viability dye (BioLegend), prepared according to the manufacturer’s instructions, at a concentration of 4 µl/ml for 30 minutes on ice. Next, samples were fixed with 4% PFA at room temperature for 30 minutes. Samples were then resuspended in 50 µl 1 x PBS, and run on an Amnis ImageStreamX Mark II. 10,000 events were collected for samples where amoebae were incubated in the absence of Jurkat cells or with Jurkat cells at a 1:5 ratio. 10,00 – 20,000 events were collected for samples with Jurkat cells at a 1:10 ratio, 10,00 - 40,000 events were collected for samples with a 1:20 ratio, and 50,000 - 80,000 events for samples with a 1:40 ratio. Data are from 6 replicates across 3 independent experiments. See Figure S2 for the analysis gating scheme.

For experiments with CD59 and CD46 knockout mutants, the serum lysis assay used was the same as described above, except only a 1:40 ratio of amoebae to Jurkat cells was used, instead of multiple different ratios. Amoebae were incubated in the absence of Jurkat cells, in the presence of Jurkat cells transfected with the px330 vector control, or with either CD59 or CD46 knockout Jurkat cell mutants at a 1:40 ratio. Cells were gated on by size and 10,000 events from the amoebae that were incubated in the absence of Jurkat cells and 100,000 events from the amoebae incubated with Jurkat cells were collected. Data are from 6 replicates across 3 independent experiments. See Figure S8a for the analysis gating scheme.

In experiments where samples were treated with phospholipase C, the assay was performed as described above with the addition of a treatment step preceding exposure to serum. Only a 1:40 ratio of amoebae to Jurkat cells was used, instead of multiple different ratios. Following coincubation of amoebae and Jurkat cells, samples were immediately placed on ice and resuspended in ice-cold M199s medium. Samples were treated with either 500 mU of Phospholipase C (PI-PLC) (Phospholipase C, Phosphatidylinositol-specific from *Bacillus cereus*; Sigma-Aldrich) prepared according to the manufacturer’s instructions, or 500 mU heat-inactivated PI-PLC. PI-PLC was heat-inactivated at 95°C for 30 minutes. PI-PLC was carried out on ice, on a rocker, in a 4°C cold room for 30 minutes. Cells were gated on by size and 10,000 events from the amoebae incubated in the absence of Jurkat cells and 100,000 events from the amoebae incubated with Jurkat cells were collected. Data are from 8 replicates across 4 independent experiments. The untreated control condition was performed 3 of 4 independent experiments and data are from 6 replicates. See Figure S8a for the analysis gating scheme.

For experiments where amoebae were incubated with increasing numbers of red blood cells, amoebae were incubated with red blood cells resuspended at 4 x 10^6^ cells/ml, 4 x 10^7^ cells/ml, and 4 x 10^8^ cells/ml for a 1:10, 1:100, and 1:1000 amoebae to red blood cell ratio. The amoeba population was gated on by size and 10,000 amoeba events were collected for samples where amoebae were incubated in the absence of red blood cells or with red blood cells at a 1:10 ratio, a 1:100 ratio. 100,000 events were collected for amoebae incubated with red blood cells at a 1:1000 ratio. Data are from 6 replicates from 3 independent experiments. See Figure S2b for the analysis gating scheme.

For C3b experiments, amoebae were labeled with CMFDA and Jurkat cells were left unlabeled. Amoebae and Jurkats were incubated together at a 1:40 amoebae: Jurkat ratio. After fixation, samples were blocked in 1 x PBST containing 5% BSA and 20% Normal Goat Serum for 30 minutes at room temperature on a rocker. Samples were then labeled with a mouse monoclonal antibody to complement components C3b and iC3b (Clone 3E7; Sigma-Aldrich) diluted 1:100 in blocking solution at 4°C overnight on a rocker. Samples were washed in 1 x PBST and labeled with an Alexa Fluor 47 AffiniPure Fab Fragment Donkey Anti-Mouse secondary antibody (Jackson ImmunoResearch Laboratories, Inc) stored in 50% glycerol (Sigma-Aldrich). The secondary antibody was diluted 1:100 in blocking solution for a final dilution of 1:200, and samples were incubated for 3 hours at room temperature on a rocker. Samples were washed in 1 x PBST, resuspended in 50 µl 1 x PBS, and run on an Amnis ImageStreamX Mark II. 10,000 events were collected in samples where amoebae were incubated in the absence of Jurkat cells and 100,000 events were collected in samples where amoebae were incubated with Jurkat cells. Data are from 6 replicates across 3 independent experiments. See Figure S3 for the analysis gating scheme.

For latex bead ingestion experiments, amoebae were left unstained and incubated with either 3×10^6^ or 1.5×10^7^ fluorescent red carboxylate-modified polystyrene 2.0 μm latex beads (Sigma-Aldrich) or incubated in the absence of latex beads. Amoebae were gated on by size and 10,000 events were collected per sample. Data are from 4 replicates across 2 independent experiments. See Figure S1 for the gating scheme used for analysis.

For experiments with amoebae exogenously expressing human CD46 or CD55, amoebae or vector control amoebae were labeled with CMFDA. Amoebae were either treated with M199s medium or exposed to human serum as described above. After gating on amoebae by size, 10,000 events were collected per sample. Data are from 8 replicates across 4 independent experiments. See Figure S8b for the gating scheme used for analysis.

### Statistical analysis

GraphPad Prism software was used to perform all statistical analyses and the means and standard deviation values are displayed on all data plots. Analyses were done using a Student’s unpaired t test (no significant difference was indicated by a P of >0.05; *, P ≤ 0.05; **, P ≤ 0.01; ***, P ≤ 0.001; ****, P ≤ 0.0001).

## ACKNOWLEDGMENTS

We thank Dr. Scott Dawson, Dr. Stephen McSorley, and the members of our laboratory for helpful discussions. All ImageStream and confocal data were acquired using shared instrumentation in the UC Davis MCB Light Microscopy Imaging Facility. We thank Dr. Michael Paddy for technical support in the use of these instruments. T.S.Y.T. was funded by the Khaira Family Experiential Learning Award through the UC Davis College of Biological Sciences. This work was funded by NIH grant AI146914 and a Pew Scholarship awarded to K.S.R.

## AUTHOR CONTRIBUTIONS

H.W.M. designed, performed, and analyzed the experiments. T.S.Y.T. created and characterized amoebae exogenously expressing complement regulatory proteins. K.S.R. conceived of the overall approach and oversaw the design and analysis of the experiments. H.W.M. and K.S.R. wrote the manuscript.

## SUPPLEMENTAL MATERIAL

### SUPPLEMENTAL FIGURE LEGENDS

**Figure S1: Gating strategy used for analysis of bead ingestion experiments**

Amoebae were gated on by brightfield area and aspect ratio during acquisition. Next, using Gradient RMS Bright Field, focused cells were gated on from total collected events. Because amoebae remained unstained in these experiments, single amoebae were then gated on using brightfield area and aspect ratio. From the single amoebae population, amoebae viability and bead ingestion were measured. Dead amoebae were gated on using fluorescence intensity of Zombie Violet dye and side scatter. Bead ingestion was quantified using the “spot count ” feature in Amnis IDEAS software and quantified spots of red bead fluorescence.

**Figure S2: Gating strategy used for analysis of experiments with increasing numbers of human cells, and for immunofluorescence analysis of amoebae exogenously expressing CD46 or CD55.**

**(A)** Gating strategy for experiments with increasing numbers of Jurkat cells, and for immunofluorescence analysis of amoebae exogenously expressing CD46 or CD55. Focused cells were gated on from total collected events, using Gradient RMS Bright Field. Single amoebae were gated using area and aspect ratio of CMFDA cytoplasm dye fluorescence. Dead amoebae were gated on using fluorescence intensity of Zombie Violet dye and side scatter. **(B)** Gating strategy for experiments with increasing numbers of red blood cells. Only amoeba events were collected for analysis and were gated on using bright field area and aspect ratio during data acquisition. Focused cells were gated on from total collected events, using Gradient RMS Bright Field. Single amoebae were gated using area and aspect ratio of CMFDA cytoplasm dye fluorescence. Dead amoebae were gated on using fluorescence intensity of Zombie Violet dye and side scatter.

**Figure S3: Gating strategy used for analysis of C3b deposition experiments.**

**(A)** Focused cells were gated on from total collected events, using Gradient RMS Bright Field. Single amoebae were gated using area and aspect ratio of CMFDA cytoplasm dye fluorescence. Dead amoebae were gated on using fluorescence intensity of Zombie Violet dye and side scatter. **(B)** Representative histograms of C3b fluorescence intensity of all single amoeba, live amoeba, or dead amoeba populations.

**Figure S4: Biotinylated human cell membrane proteins are detected on the surface of amoebae prior to fixation.**

The surface of human Jurkat cells was biotinylated prior to incubation with amoebae. Following incubation, samples were placed on ice to halt membrane turnover and fluorescently-conjugated streptavidin was used to detect biotinylated proteins on the surface on both human cells and amoebae (magenta) prior to fixation. DNA was labeled with DAPI nucleic acid stain following fixation. Arrows indicate transferred patches of human proteins on the surface of amoebae. **(A)** Amoebae and biotinylated human cells were incubated together for 2 minutes. **(B)** A closeup image of an amoebae from the panel shown in A with transferred human proteins on its surface. **(C)** Amoebae and human cells were incubated together for 5 minutes. Amoebic autofluorescence is shown in green. **(D)** 3-D reconstruction of Z stacks taken from the data in C. **(E-F)** Human cells and amoebae were incubated together for 5 minutes. Amoebae cytoplasm was labeled with CMFDA dye (green), and the nuclei of cells were left unstained. Data were analyzed by confocal microscopy. Images in A-F are representative of data collected from 4 independent experiments with incubation times of 2-5 minutes.

**Figure S5: Gating strategy used for analysis of CD59 displayed on amoebae after 5 minutes and 1 hour of trogocytosis.**

A masking strategy was developed to quantify only fluorescence of CD59 present on the amoebae, and not on extracellular human cells. **(A)** Focused cells were gated on from total collected events, using Gradient RMS Bright Field. Single amoebae were gated using area and aspect ratio of CMFDA cytoplasm dye fluorescence. Next, fluorescence intensity of CD59 inside of the masked area was measured. **(B)** Representative images of bright field, amoeba cytoplasm, human cell nuclei, and CD59 fluorescence with the masked area (turquoise) applied as an overlay.

**Figure S6: Amoebae acquire and display the complement regulatory protein CD46 from human cells.**

Amoebae were incubated in the absence of Jurkat T cells, or were allowed to perform trogocytosis on human Jurkat cells for 5 minutes. Human cell nuclei were pre-labeled with Hoechst (blue), and amoebae were pre-labeled with CMFDA (green). Human CD46 (red) was detected on the amoebae surface by monoclonal antibody staining. (A) Representative images from amoebae that were incubated in the absence of Jurkat cells. (B) Representative images of amoebae that performed trogocytosis on human Jurkat cells for 5 minutes. Arrows indicate patches of displayed CD46 on the amoeba surface. Data were analyzed by imaging flow cytometry and are from 2 replicates and 1 independent experiment.

**Figure S7: Removal of GPI-anchored surface proteins using phosphatidylinositol-specific phospholipase C.**

Amoebae were labeled with CMFDA cytoplasm dye and incubated in the absence of Jurkat cells or with Jurkat cells. Human cells were labeled with DiD membrane dye. Samples were then treated with phosphatidylinositol-specific phospholipase C (PI-PLC) to remove GPI-anchored proteins, or heat-inactivated phosphatidylinositol-specific phospholipase C (HI-PI-PLC) as a control. Following exposure to human serum, amoeba death was assessed with Zombie Violet viability dye and ingested human cell material was determined by quantifying mean fluorescence intensity (MFI) of DiD present on amoebae. **(A)** Normalized death of amoebae. **(B)** Normalized mean fluorescence intensity of DiD on amoebae. Data were analyzed by imaging flow cytometry and are from 8 replicates across 4 independent experiments. The untreated control condition was performed 3 of 4 independent experiments and data are from 6 replicates.

**Figure S8: Gating strategy used for analysis of experiments with CRISPR knockout mutants, PI-PLC treatment experiments, amoebae exogenously expressing CD46 or CD55.**

Only cell events were collected for analysis (to minimize collection of debris) and were gated on using bright field area and aspect ratio during data acquisition. Focused cells were gated on from total collected events, using Gradient RMS Bright Field. Single amoebae were gated using intensity of and aspect ratio of CMFDA cytoplasm dye fluorescence. **(A)** In CRISPR knockout mutant and PI-PLC treatment experiments, amoebae were gated on a second time using CMFDA fluorescence intensity and side scatter to eliminate remaining clumps of human cells from the analysis. **(B)** In experiments with amoebae exogenously expressing CD46 or CD55, the second amoebae gate was not used. Finally, dead amoebae were gated on using fluorescence intensity of Zombie Violet dye and side scatter.

**Figure S9: Sequencing analysis of Jurkat CRISPR/Cas9 mutants.**

Chromatograms from Sanger sequencing analysis of Jurkat T cell CRISPR/Cas9 mutants. Chromatograms from CD59 mutants **(A)** and CD46 mutants **(B)** show that the gene sequence of the mutants is different from the vector control cells downstream of the gRNA/PAM sites.

