## Supplementary figures and images for "*Entamoeba histolytica* develops resistance to complement deposition and lysis after acquisition of human complement regulatory proteins through trogocytosis"

### Figure S1

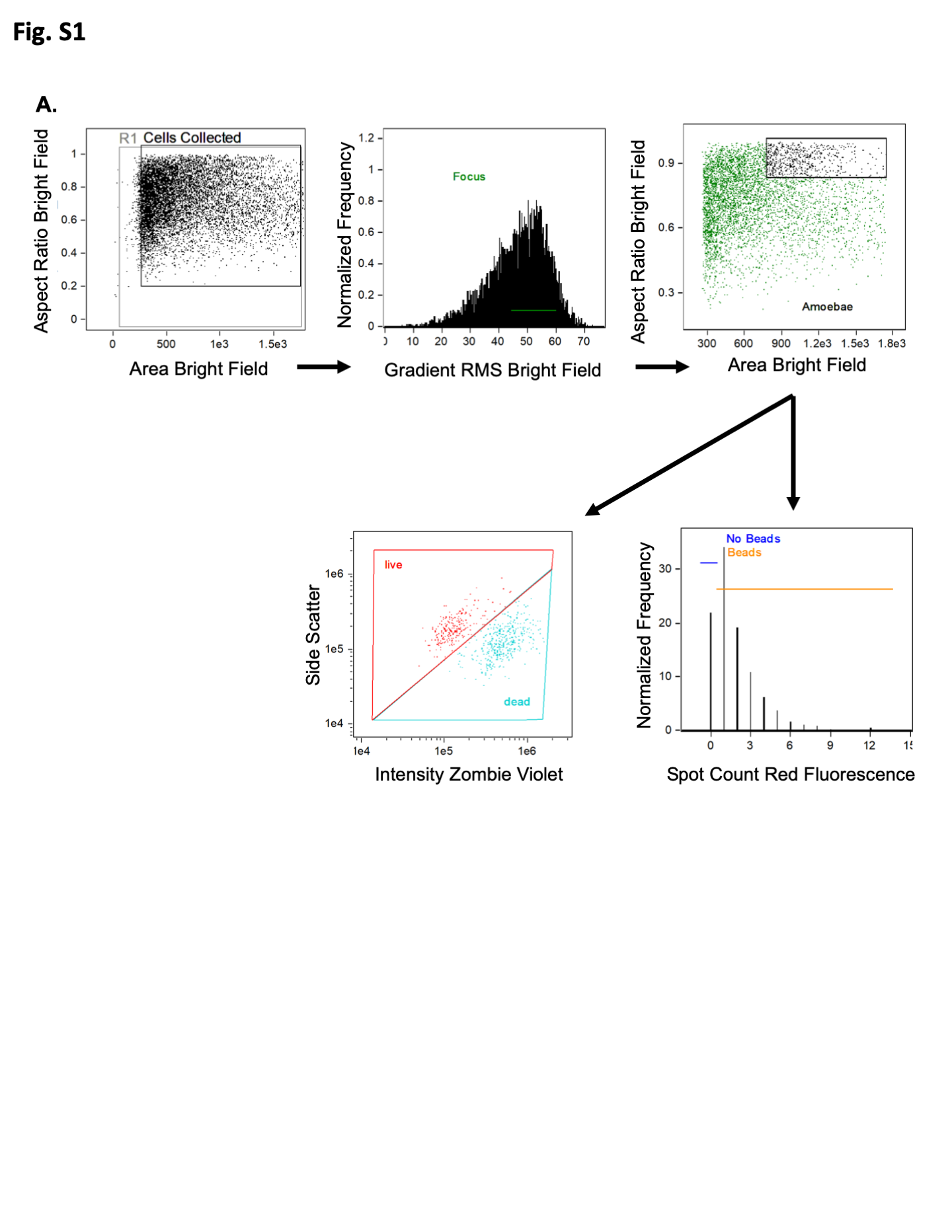

### Figure S2

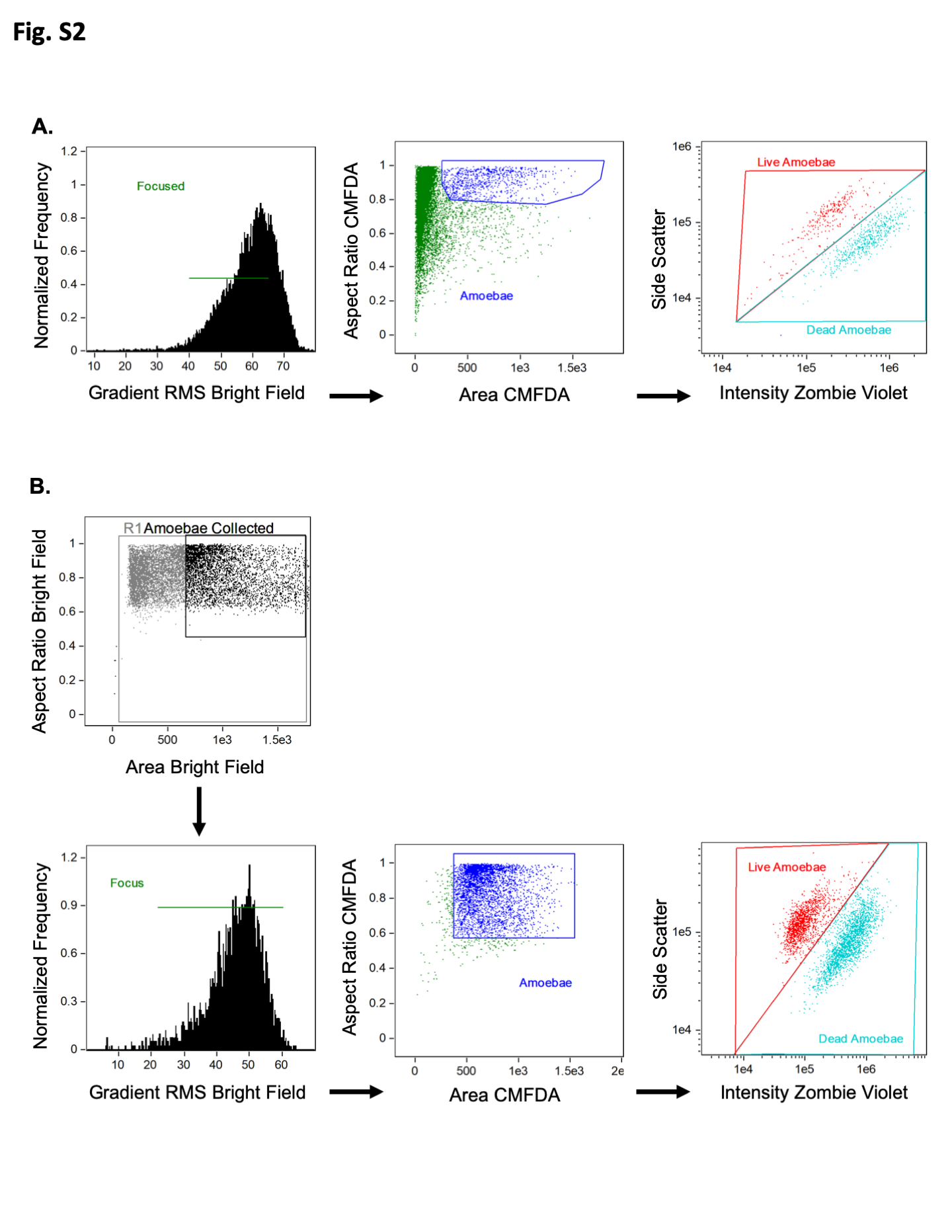

### Figure S3

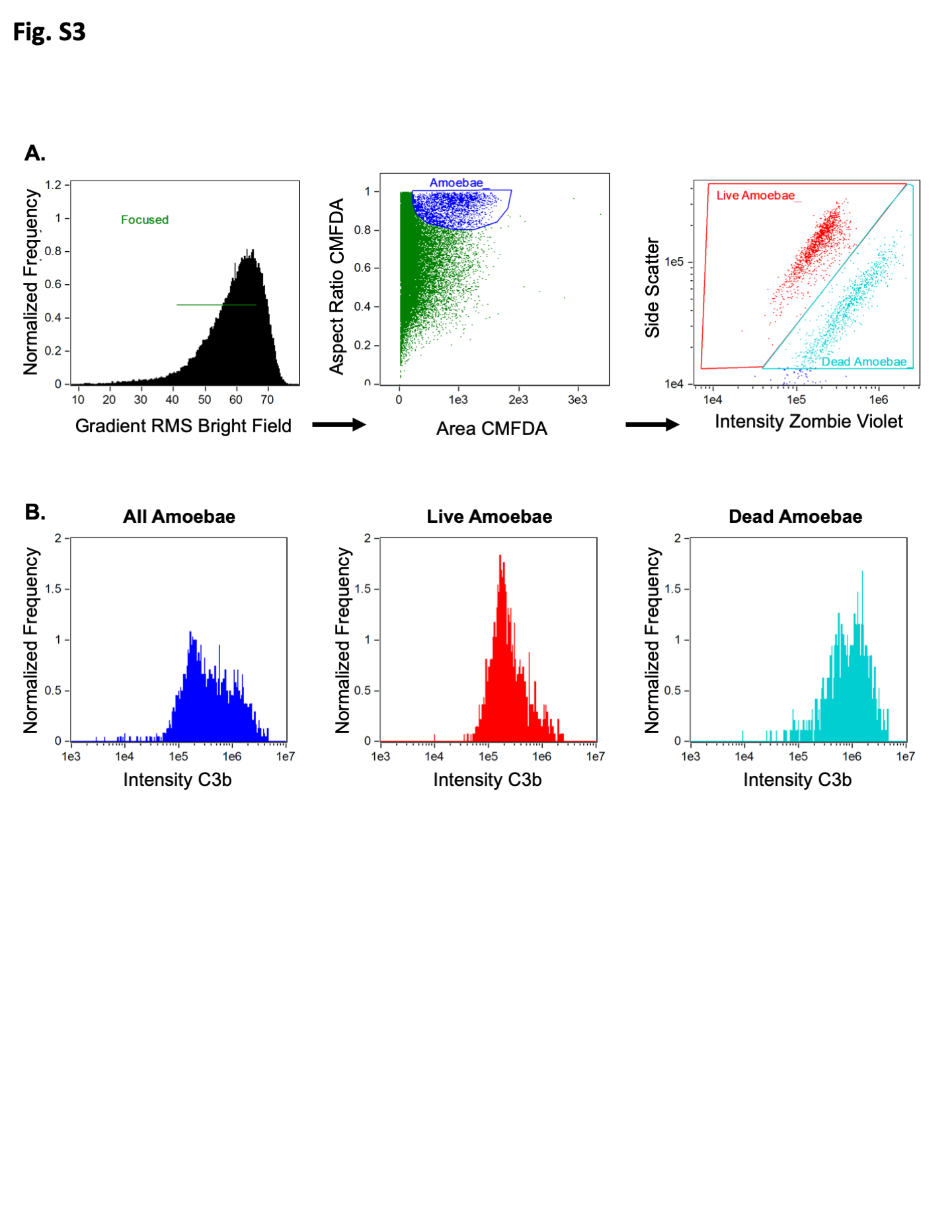

### Figure S4

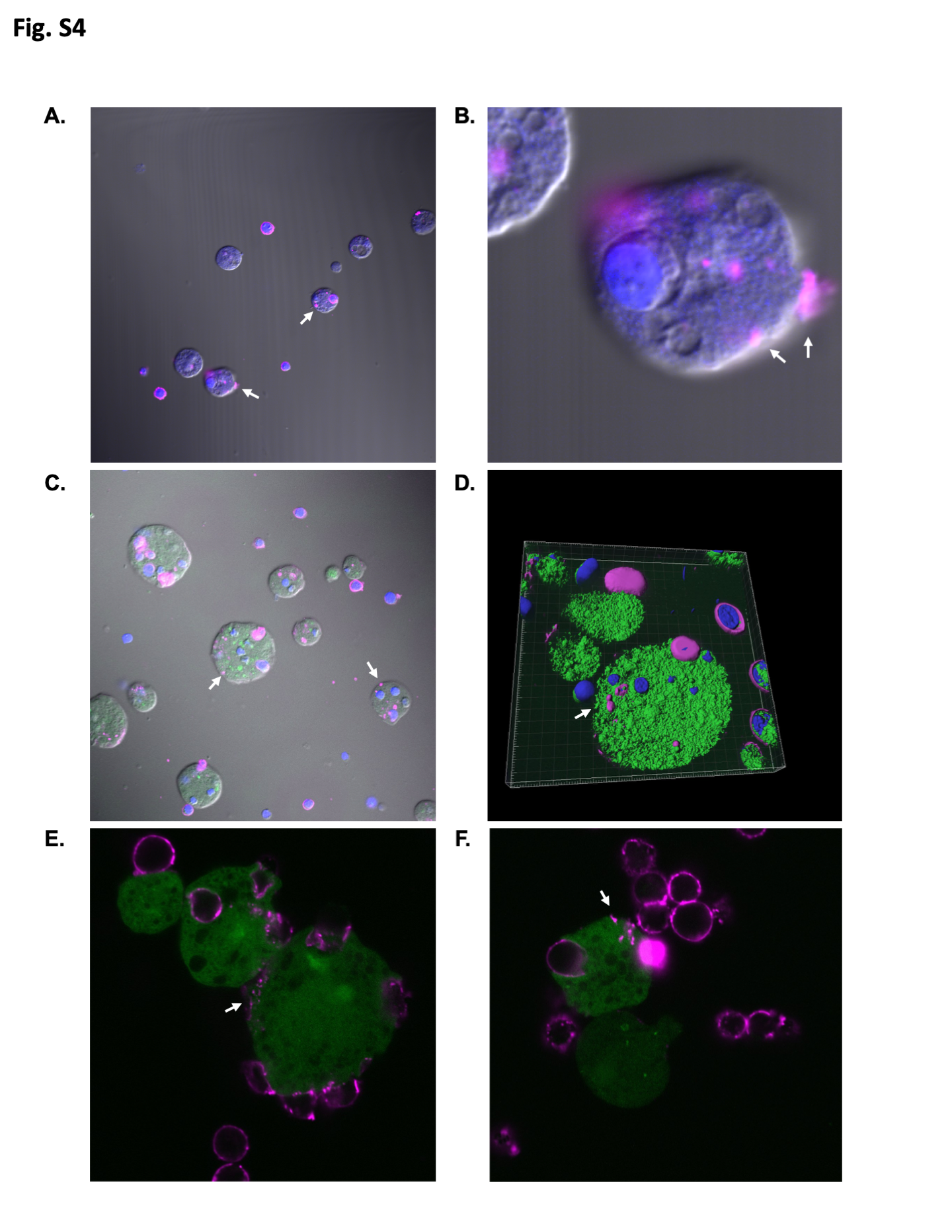

### Figure S5

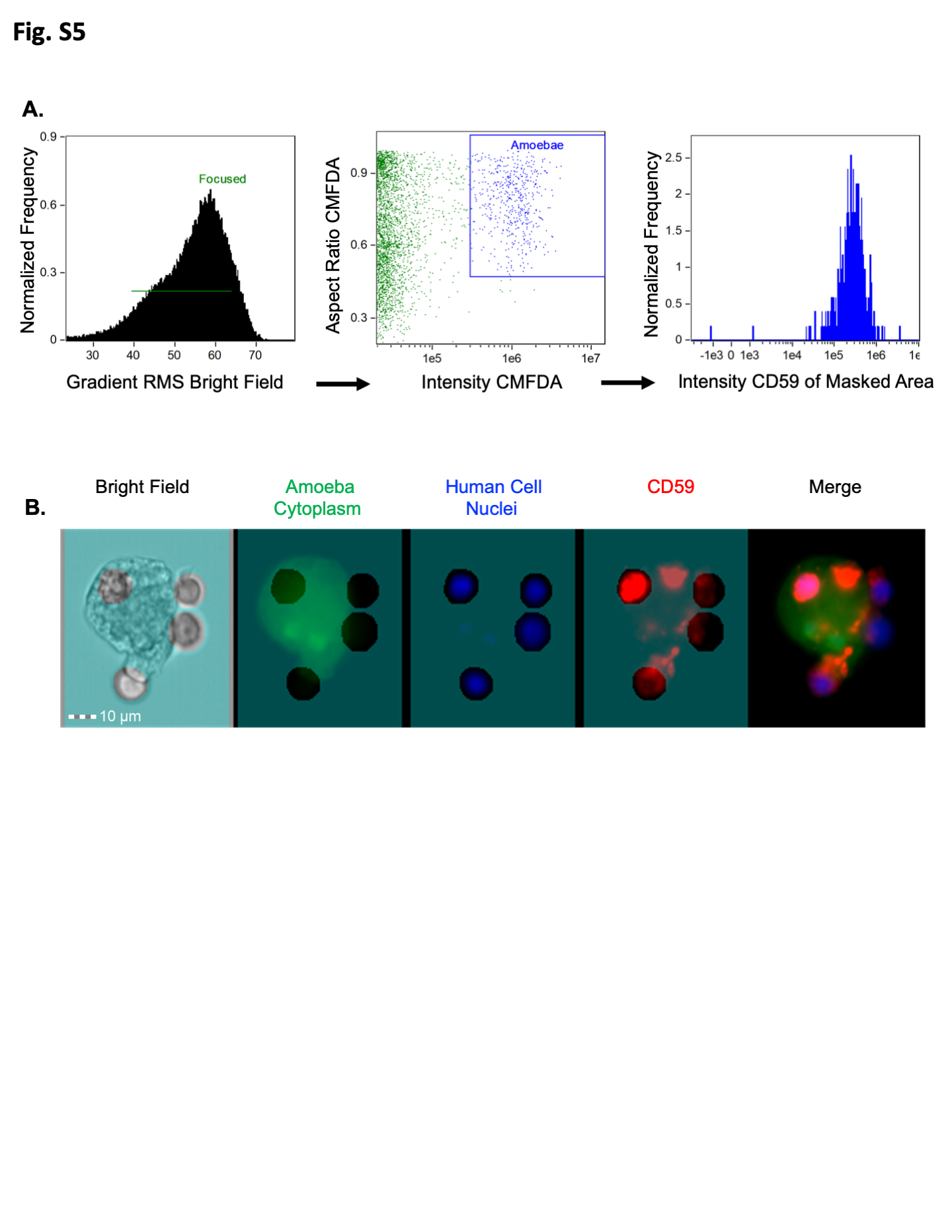

### Figure S6

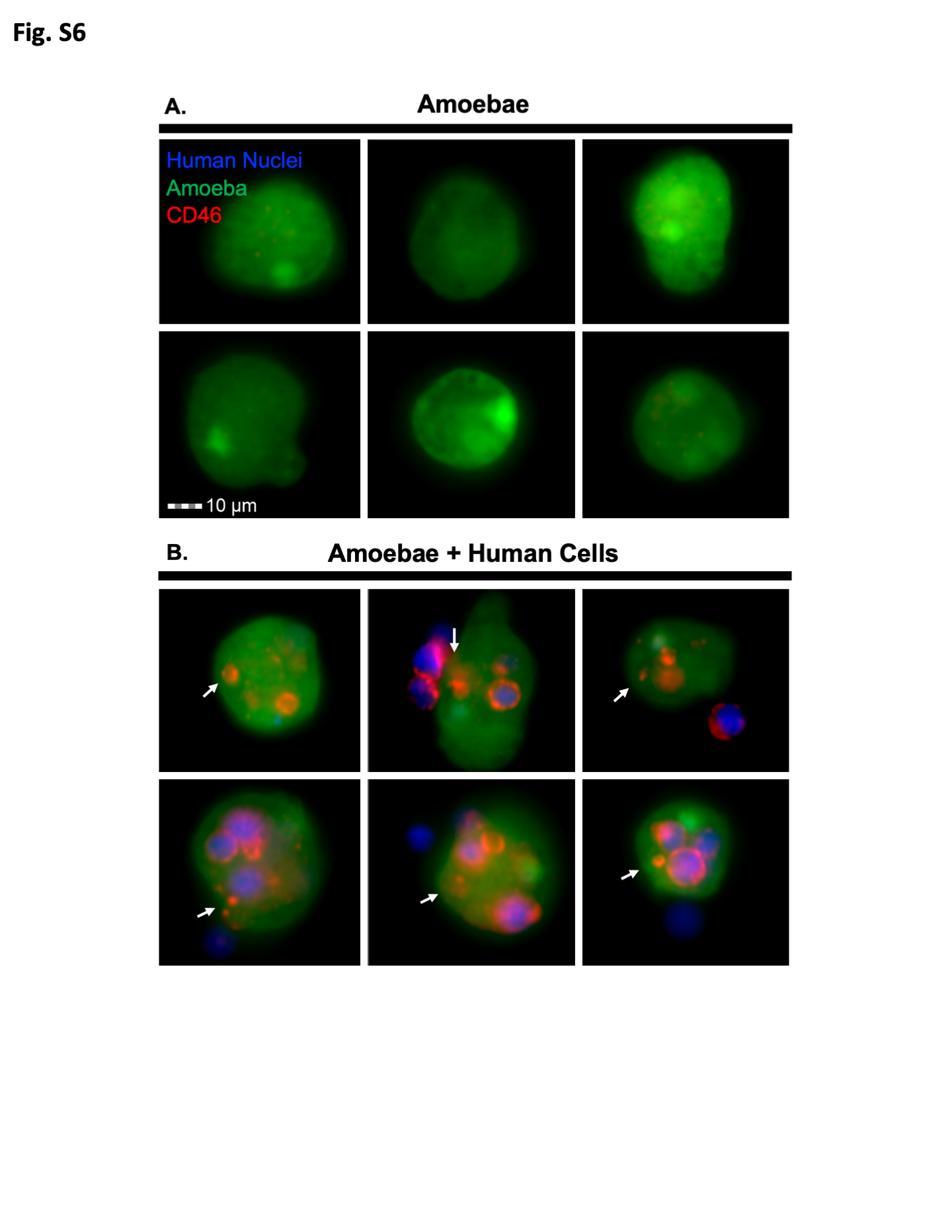

### Figure S7

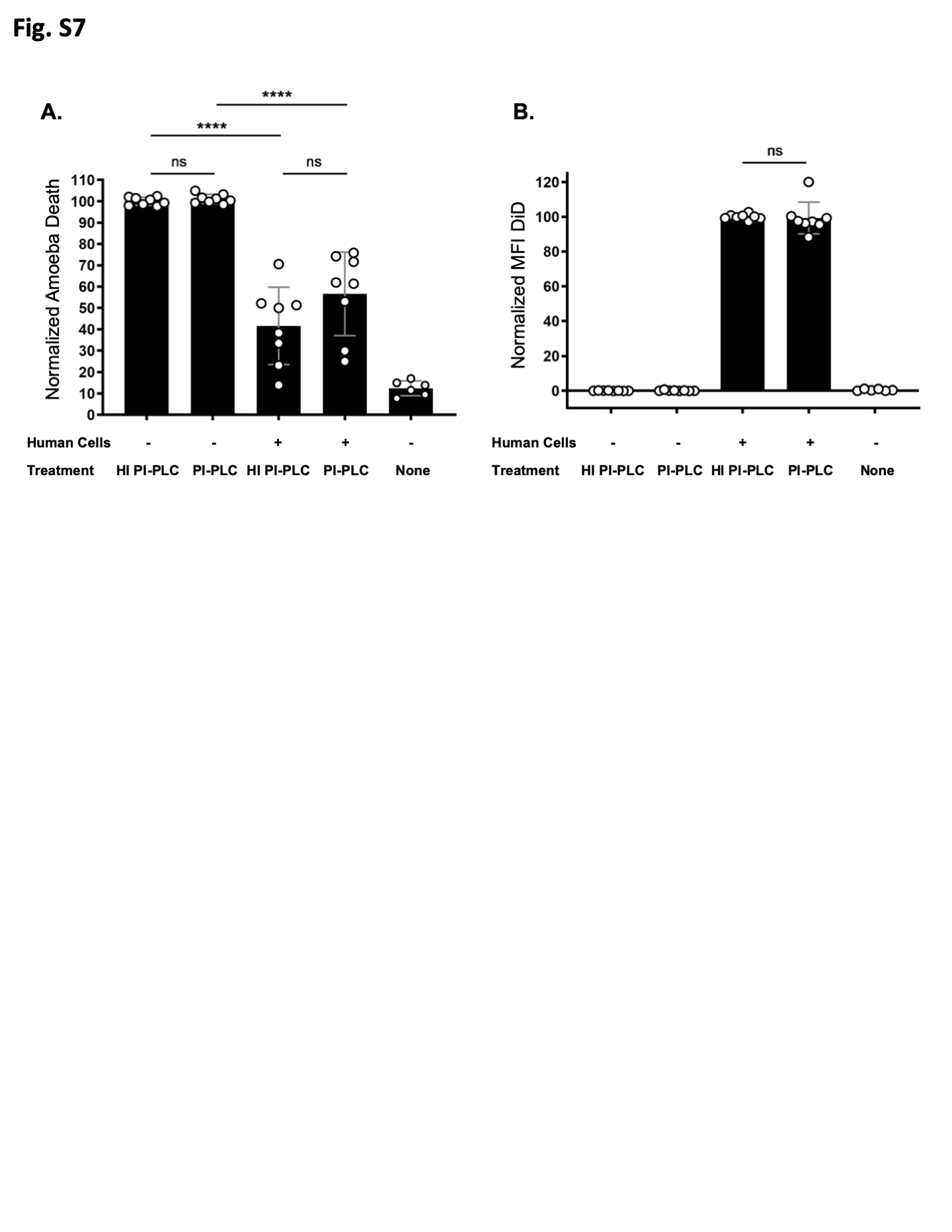

### Figure S8

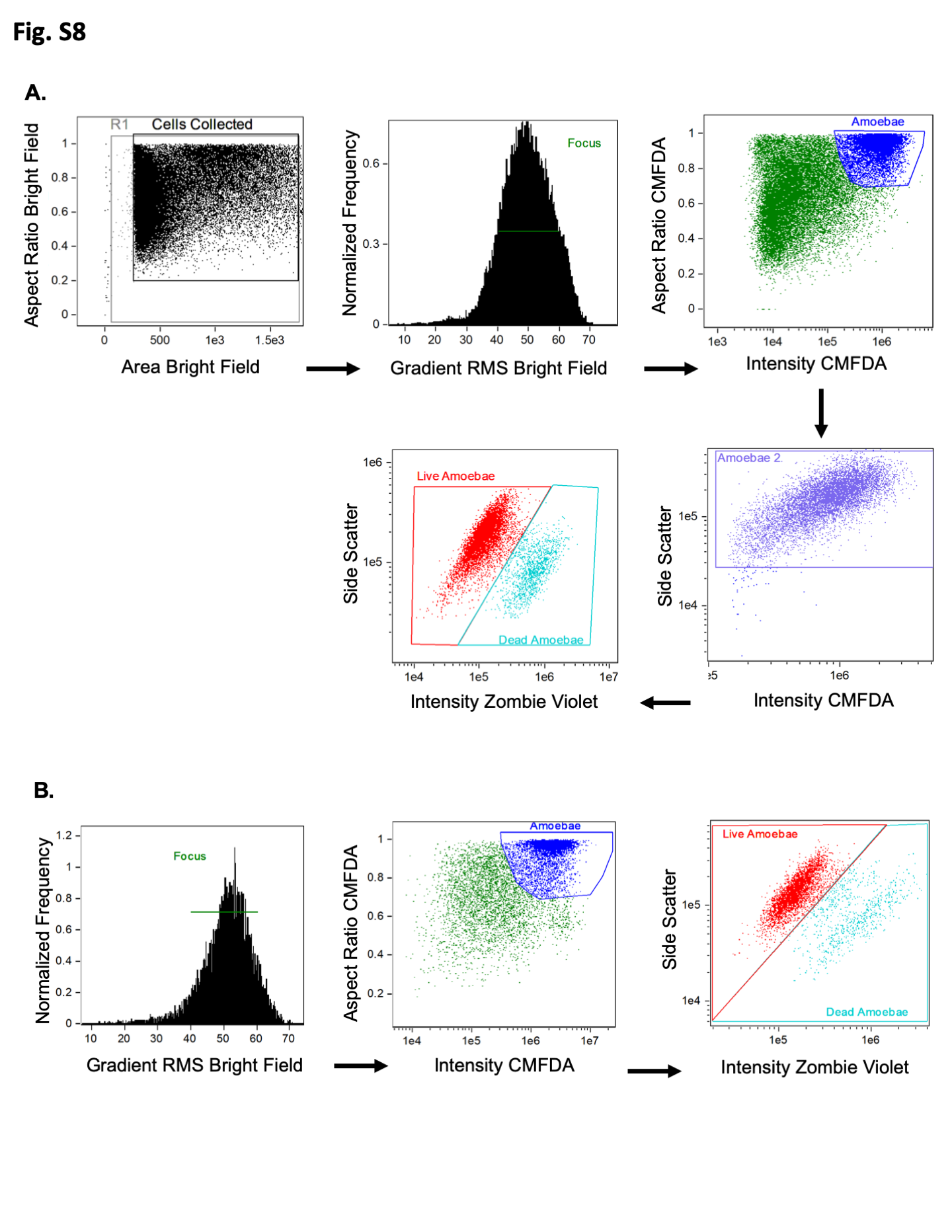

### Figure S9

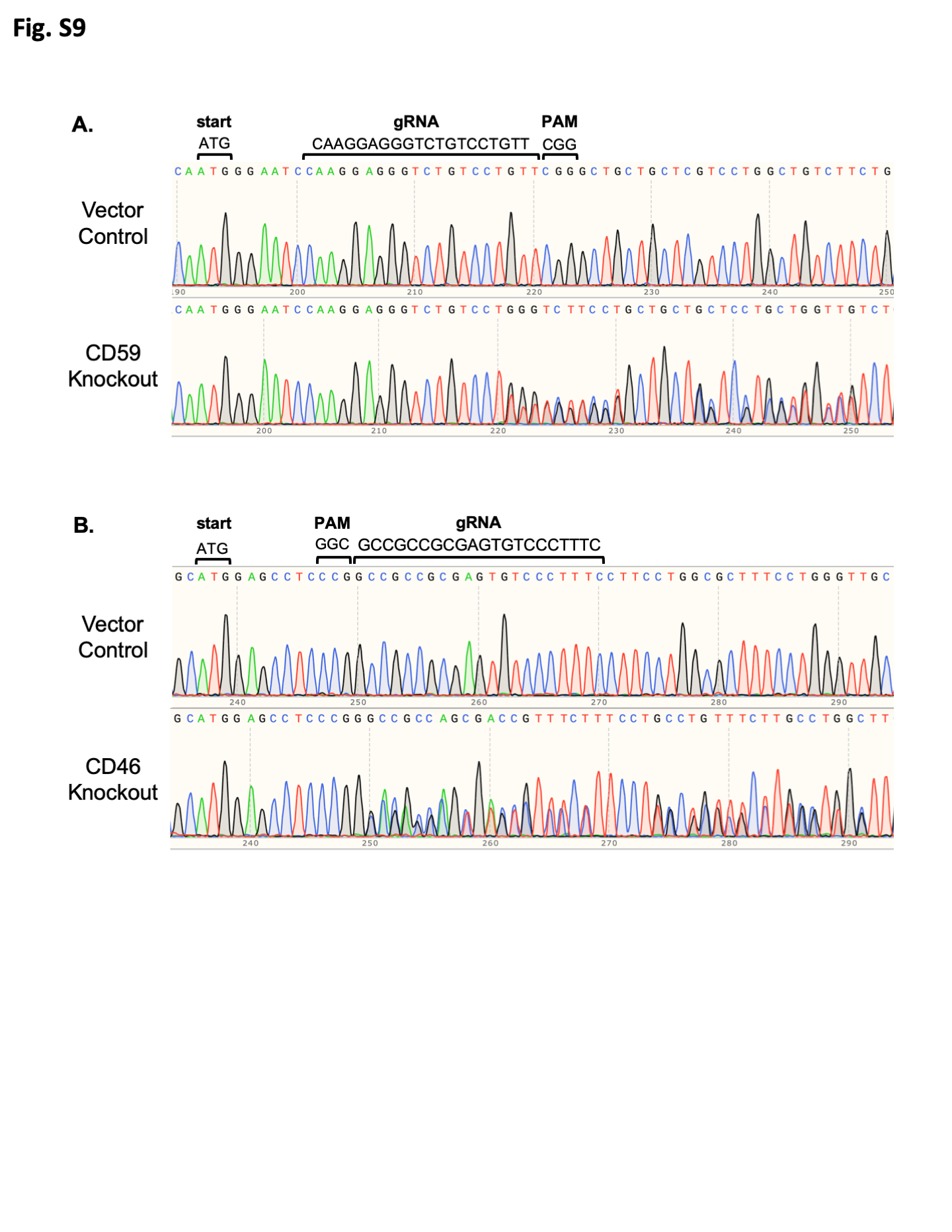
